# Setting priorities for the acquisition of primary plant occurrence data

**DOI:** 10.1101/2025.11.11.683036

**Authors:** Nadia Bystriakova, Pablo Alvarez Hendrigo De Melo, Alexandre Antonelli, Steven Bachman, Gemma Bramley, Matilda Brown, Gloria Cespedes, Martin Cheek, Iain Darbyshire, Sebsebe Demissew, Juana DeEgea, Andre Erst, Felix Forest, Ib Friis, Long-Fei Fu, Alfredo Fuentes, Rajib Gogoi, Laura Jennings, Carel C.H. Jongkind, Bente Klitgaard, Isabel Larridon, Eve Lucas, Carla Maldonado, Maira Martinez, Justin Moat, Eimear Nic Lughadha, Carlos Reynel, Himmah Rustiami, Daniel Santamaria Aguilar, Sebastian Tello, Liam Trethowan, Timothy M.A. Utteridge, Bat (Maria) Vorontsova, Yi-Gang Wei, Tom Wells, Alexandre K. Monro

## Abstract

**Aim:** Effective implementation of the Global Biodiversity Framework and Global Strategy for Plant Conservation depends on accurate species distribution data. Current vascular plant distribution data, while crucial for understanding terrestrial ecosystems, is often sparse and biased and requires significant expansion. This study developed a scalable approach to prioritize areas for plant occurrence data acquisition, adaptable to national priorities and providing a framework for botanical institutions to coordinate efforts and allocate resources.

**Location:** Global.

**Methods:** Using a Technique for Order Preference by Similarity to Ideal Solution (TOPSIS) analysis, we prioritized areas based on: (a) ecosystem service value; (b) floristic value threatened by climate or land-use change; and (c) uncertainty in species richness estimates, stratified by biome and region. Regional prioritization maps for Africa & Madagascar; East, South and Southeast Asia; Siberia and the Russian Far East; South America; and North & Central America were reviewed by botanical experts for validation. Scalability was assessed by comparing regional and global analyses.

**Results:** Data-driven priority maps, divided into tree-dominated and grassland/deforested areas, largely received expert support. High similarity between global and regional maps demonstrated scalability.

**Main conclusions:** Our approach provides a framework for supporting national implementation of the Global Biodiversity Framework. Variables and their weights can be tailored to national or local needs. The method’s flexibility and adaptability extend to other taxonomic groups and objectives, such as protected area selection By prioritizing data acquisition, whether field-based or digital, this research promotes the efficient use of resources. A key advantage of this approach is its capacity to systematically translate expert opinion into explicit and quantitative criteria, which in turn facilitates clear communication with policymakers and funders.

## Introduction

The current human-driven mass extinction event (Barnosky et al., 2011) poses a significant threat to the survival of many species with which we share the Earth (Ceballos et al., 2015) and the ecosystem services on which we rely (Díaz et al., 2019). In response, the international community has developed the 2050 Vision for Biodiversity, which includes the Global Biodiversity Framework (GBF; Convention on Biological Diversity, 2022; GBIF, 2025). For plants, a new Global Strategy for Plant Conservation (GSPC) comprises 21 actions to complement the GBF (Convention on Biological Diversity, 2012), eight of which are dependent upon an accurate knowledge of plant occurrences. Both the GBF and GSPC rely on accurate evidence of species and their distributions, and of areas and their species assemblages. For example, achieving Target 3 (Conserve 30% of Land, Waters, and Seas), requires the identification and monitoring of areas of particular importance for biodiversity (Convention on Biological Diversity, 2022). Target 21 emphasizes the need for policymakers, practitioners and the public to access the best available data. Furthermore, the effectiveness of other key targets (e.g., Targets 1, 2, 4, and 14) also hinges on the availability of high-quality data to guide conservation actions.

Consideration of coverage and resolution are vital in determining whether occurrence data are sufficient to support effective implementation and predictions of species distributions (Moudrý et al., 2024). For example, the data needs to be capable of supporting the accurate predictions of species distribution below the national scale. Implementation of the CBD typically occurs at the national or subnational level. Identifications and spatial coordinates also need to be verifiable, with preserved specimen data being the gold-standard for plants, as for Red Listing (Nic Lughadha et al., 2019).

The diversity of most organisms that underpin terrestrial ecosystem function —such as invertebrates, fungi, and microbes, and plants —remains insufficiently documented, making it difficult to accurately infer species distributions for those groups (IPBES, 2019). Consequently, they are currently less integrated in the implementation of the (GBF) than groups such as mammals, birds and amphibians. This might be less of an issue for vascular plants as their diversity is better documented (Cheek et al., 2020). Primary data on vascular plant species occurrences, however, are both sparse in terms of the number of vouchered specimens and very biased spatially, taxonomically and temporally (Feeley & Silman, 2011; Feng et al., 2022; Hortal et al., 2015; Meyer et al., 2016; Nelson & Ellis, 2019; Ramírez-Barahona et al., 2023) resulting in intrinsic limitations to its application to studying plant diversity and distribution (Button & Borzée, 2024; Bystriakova et al., 2012; Hortal et al., 2015). The proportion of digital occurrence data that is verifiable and including species identifications is also ever-decreasing (Troudet et al., 2018) as the generation of digital citizen-science observations and plot observation data increases.

Data scientists rely on species distribution models (SDMs) to predict species’ occurrences and investigate diversity patterns (Cai et al., 2023; Jung et al., 2021; Ondo et al., 2024; Sabatini et al., 2022). Models that incorporate primary occurrence data, however, are rarely externally validated (Alves Martins et al., 2024; Cai et al., 2023; Moura & Jetz, 2021; Newbold et al. 2010; Ondo et al., 2024), and risk generating distorted results. A significant challenge with the use of SDMs is the absence of sufficient training occurrence data (Moudrý et al., 2024) and strong sample bias (Feeley & Silman, 2011). To generate a reliable prediction, SDMs require 20 (Mateo et al., 2010; exceptionally 5 data points for range-restricted species (Hernandez et al. 2005)) to 200 (Hanberry et al., 2012) occurrences from across a species’ range. Estimates of around 400,000 vascular plant species (Antonelli et al. 2023) suggest that 8 to 80 million rigorously sampled occurrences could enable reliable distribution predictions. Although GBIF holds approximately 113 million vascular plant occurrences (2024; i.e. preserved specimens, continuously updated), once parsed and those lacking spatial coordinates or species identifications are excluded (de Melo et al. 2024) these represent 23 million unique occurrences, the distribution of which is highly uneven (Fig.1), Enquist et al. (2019) finding that about one-third of vascular plant species have five or fewer records. There also remain important biological collections whose occurrence records have not been digitised. For example, the Komarov Institute, Russia.

**Fig. 1.**
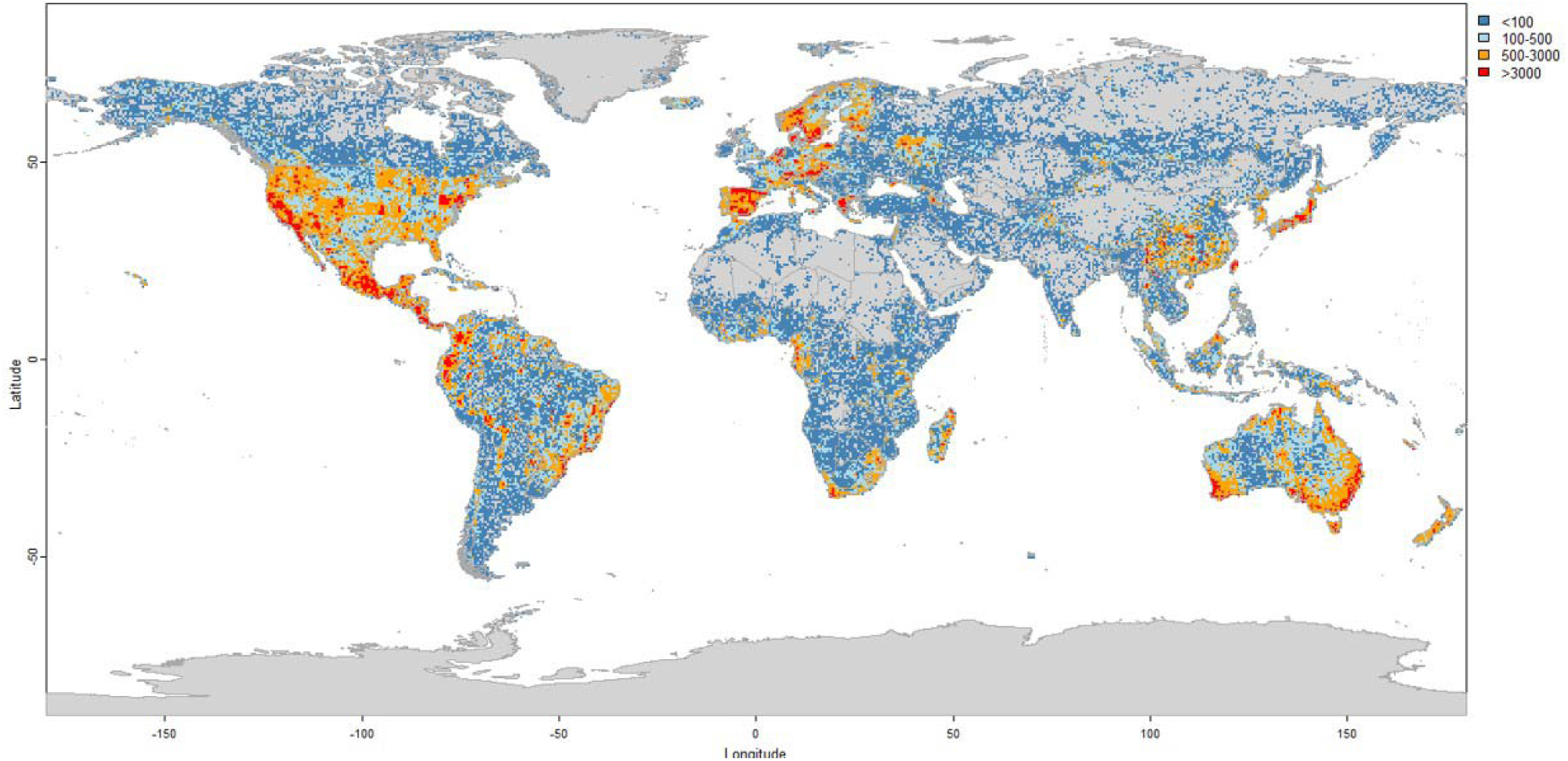
Number of parsed GBIF preserved-specimen occurrence records across the globe, aggregated at a 50×50 km resolution. Data comprises records that have both spatial coordinates and a species determination. GBIF occurrence records were parsed using parseGBIF work package (de Melo et al. 2024). Total of 23 million parsed records. Records available at https://doi.org/10.6084/m9.figshare.25849084.

In summary, our current knowledge of plant occurrences is insufficient to effectively prioritise areas with confidence on a global scale to estimate species richness, promote scientific discovery, or Red List assessments of extinction threat (Pimm et al., 2014). Additional primary data is urgently needed, not only to meet the GBF and GSPC targets by 2030 but also to support broader biodiversity-positive actions (Antonelli et al., 2023; IPBES, 2019) and more reliable SDMs (Feeley & Silman, 2011). Such additional data can be generated or mobilised in several ways: 1) through additional targeted ground inventories, 2) through the integration of the extensive network of vegetation plots and preserved specimen collections 3) the extraction of meta-data from herbarium vouchers currently unusable because they lack either spatial coordinates or a species identification, and 4) the digitisation of specimens in herbarium collections. Generating additional primary data would not only address gaps in plant distribution data but also improve the models used to predict species distributions and inform Red List assessments (Rocchetti et al., 2021). However, ongoing data collection does not efficiently mitigate existing biases, often re-enforcing them (Di Cecco et al., 2021; <u>Farooq et al., 2021</u>; Geldmann et al., 2016; Geurts et al., 2023) as collector behaviour, driven primarily by accessibility and previous knowledge, remains unchanged. In addition, the botanical community has long lacked a strategy to optimise data acquisition efforts to address these biases (Antonelli et al., 2024).We propose a new approach to prioritizing the acquisition of additional primary data, with the aim of countering existing biases and prioritising those areas most at risk, underpinning the creation of a primary data set that will support measures enabling us to reach the global vision of a world living in harmony with nature by 2050.

The criteria by which areas are prioritised as foci for data acquisition should be determined by the needs, ambitions and resources of end users. Broadly applicable criteria would include the mitigation of sample bias (e.g. with respect to habitats and or climate regimes), the degree of urgency related to threats to species and areas, and the provision of ecosystem services. Locally applicable criteria could include the need to prioritize globally common but nationally scarce biomes, or areas rich in species of importance for livelihoods.

Estimates of sampling effort (Lobo et al., 2018) and bias (Zizka et al., 2021) are commonly employed. To prioritise areas for the acquisition of additional primary species occurrence data, with the aim of mitigating existing biases. For example, survey completeness and spatial bias were estimated to identify priority areas for new data acquisition in Bignoniaceae (Narváez-Gómez et al., 2021). These estimates have also been combined with taxonomic and geographic shortfalls analyses to identify global biodiversity ‘darkspots’ (Ondo et al., 2024). Chanachai et al. (Chanachai et al., 2024) “identified forested areas for future botanical exploration”, by examining the spatial overlap between inventory completeness, remaining natural habitat and protected areas and degree of forest modification.

In this study we aimed to use a broad range of criteria, encompassing biodiversity and ecosystem services potential, threats to the areas or taxa that those areas encompass, and uncertainty surrounding current floristic diversity estimates (Table 1), to assemble an occurrence data set that can most effectively support the GBF and GSPC. We sought to mitigate methodological issues identified by Button & Borzee (2024) and Sabatini et al (2022) by avoiding a reliance on direct measures of plant species richness and by compartmentalising the data by geographical region and biome (i.e. tree-dominated or grassland/deforested).

**Table 1.** Environmental variables (layers) selected for future sampling effort prioritization.

| Criterion | Variable | Citation |
| --- | --- | --- |
| (a) | Areas of global importance for conserving terrestrial biodiversity, carbon, and water | Jung et al. (Jung et al., 2021) |
| (a) | Climate stability from Pliocene (3.3 Ma) to the present (species richness driver) | Herrando-Moraira et al. (Herrando-Moraira et al., 2022) |
| (a) | Terrain roughness (species richness driver) | Wilson et al. (Wilson et al., 2007) |
| (b) | Global human modification of terrestrial systems | Kennedy et al. (Kennedy et al., 2020) |
| (b) | Climate stability from present to 2100 | Herrando-Moraira et al. (Herrando-Moraira et al., 2022) |
| (b) | Proportion of observed species predicted to be threatened | Bachman et al. (Bachman et al., 2024) |
| (c) | Sampling bias | Zizka et al. (Zizka et al., 2021) |
| (c) | Normalized difference between recorded and expected taxon richness | Cai et al. (Cai et al., 2023)<br>de Melo (de Melo et al., 2024) |

The objective of this study was to determine whether a multi-criteria approach, integrating environmental, floristic, and ecosystem variables, can be used to effectively identify and rank priority areas for future botanical data acquisition. To answer this, we used a multi-criteria-based decision-making Technique for Order Preference by Similarity to Ideal Solution (TOPSIS), a mathematical tool for solving multicriteria decision problems (Hwang Ching-Lai & Yoon, 1981). A set of priority areas identified by TOPSIS were subsequently revised by field experts.

## Methods

The complete workflow developed here is presented in Fig. 2, and the major steps are described in more detail below. In addition to a global scale analysis, we used Africa and Madagascar; East, South and Southeast Asia, Siberia and the Russian Far East, and the Americas as case studies to validate the applicability of the workflow across different regions selected because they include many of the biodiversity hotspots identified by previous authors (Davis et al., 1997; Myers et al., 2000). The analysis was conducted separately for each region to account for their distinct ecological and geographical characteristics.

**Fig. 2.**
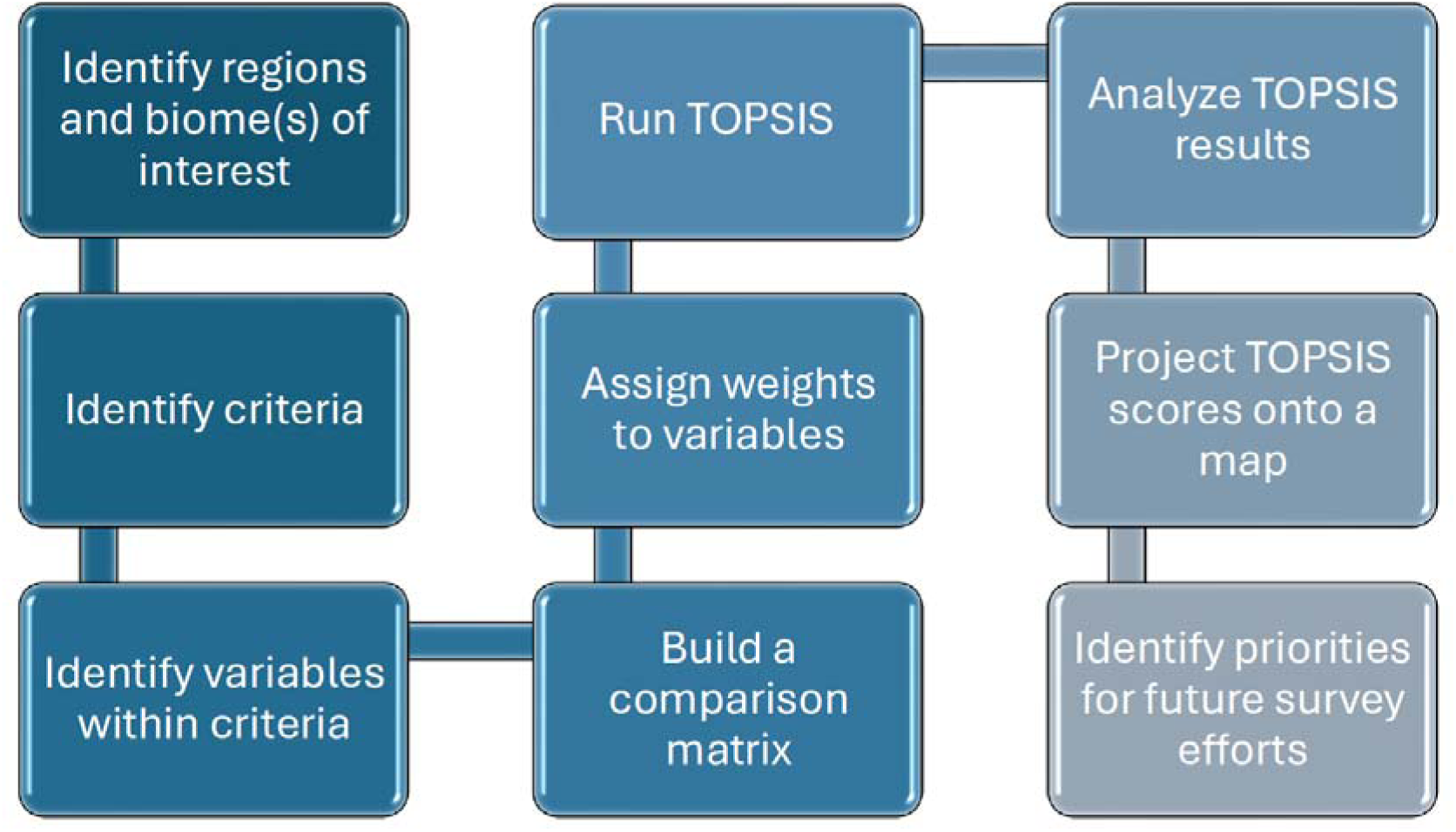
Workflow used to prioritize areas for the acquisition of additional primary occurrence data.

### Criteria and variable selection

We determined that areas for new data acquisition should satisfy at least one of the following broadly defined criteria: (a) high value of ecosystem services, including biodiversity; (b) high threats to plant diversity due to climate or land use change; c) high uncertainty of species composition estimates. To facilitate area selection based on these criteria, we used eight environmental variables available at the global scale (Table 1, Supplement 1). The variable choice was limited by availability of floristic data at the global scale with a resolution higher or equal to that selected for the analysis (50 x 50 km).

For **criterion (a)**, which includes carbon and water resource together with a measure of biodiversity, we identified three environmental variables linked to high species richness and biodiversity. The first variable is a result of a global study that ranked areas based on their importance for terrestrial biodiversity conservation and nature’s contributions to carbon storage and water (Jung et al., 2021). Such areas require the estimation of species richness and turnover and therefore should be prioritized for additional data acquisition.

The second variable representing climate stability from Pliocene to the present was selected to incorporate the notion of climatically stable areas (Colville et al., 2020; Cowling et al., 2015). These areas of climate stability are considered a priority for future sampling efforts as they can provide insights into the evolutionary processes that shaped modern flora (Herrando-Moraira et al., 2022).

Evidence suggests a strong relationship between habitat heterogeneity and species diversity (Antonneli, 2018; Báldi, 2008). In this study, terrain roughness, defined as the difference between the maximum and the minimum value of elevation of a cell and its eight surrounding cells, was used as a proxy for habitat diversity (Wilson et al., 2007). This metric was selected following a comparison with the terrain roughness index (TRI, Wilson et al., 2007) in which we observed TRI to have less discriminatory power than elevation range at a cell size of 50 x 50 km and so opted for the former.

The drivers of biodiversity loss are well known and extensively documented (IPBES, 2019). Land use remains the greatest contributor to biodiversity loss, closely followed by climate change (Boakes et al., 2024). To incorporate **criterion (b)**, we focused on three direct drivers that could be easily measured and visualized, and for which digital maps were freely available: global human modification of terrestrial systems, future climate stability, and extinction threat.

The Global Human Modification variable represents the degree of human modification of all terrestrial lands worldwide (Kennedy et al., 2020). The variable combines five major categories of human pressures: 1) human settlement (population density, built-up areas), 2) agriculture (cropland, livestock), 3) transportation (major roads, minor roads, two-tracks, and railroads), 4) mining and energy production (mining/industrial areas, oil wells and wind turbines), and 5) electrical infrastructure (powerlines and night-time lights). The dataset was downloaded from https://www.earthdata.nasa.gov/.

Climate change is a key driver of biodiversity loss on the global scale (Vera et al., 2023). For future data acquisition, we prioritized areas which are expected to be strongly affected by climate change over those with low climatic fluctuations. We used a map of Climate Stability Index (CSI) values for the future conditions under the Shared Socioeconomic Pathway SSP1-2.6, from the present (1970–2000) to future (2100); this scenario assumes the lowest emission level (Herrando-Moraira et al., 2022) and has been demonstrated to be the most realistic (Garner & Pearson, 2023).

To estimate threatened species richness in 50×50 km grid cells, we linked two datasets: a list of vascular plant taxa identified as “threatened” based on IUCN Red List assessments and/or machine learning predictions (Bachman et al., 2024) and a dataset of all vascular plant occurrences from GBIF (www.gbif.org), verified using the ParseGBIF workflow (de Melo et al., 2024a, de Melo et al., 2024b). We retrieved distribution data for 111,605 unique taxa estimated to be threatened. For future data acquisition, we prioritized grid cells with the largest number of threatened species.

To address **criterion (c)**, we used two measures of sample effort, sampling bias and the difference between estimated and observed taxon richness. Sampling bias was calculated at the global scale using distribution data of all vascular plants retrieved from GBIF (www.gbif.org). Over 200 preserved specimen datasets were downloaded by family and are available on Figshare (Monro & de Melo 2024) and verified by ParseGBIF workflow (de Melo et al., 2024a, de Melo et al., 2024b), with the help of the R package “sampbias” (Zizka et al., 2021) and “parseGBIF” (https://github.com/pablopains/parseGBIF).

As a proxy for estimated vascular plant species richness, we used plant diversity maps in raster format at a resolution of 30 arc seconds (ca 1 km) produced by GIFT (Cai et al., 2023). The observed species richness was represented by the distribution data of all vascular plants accumulated by GBIF (www.gbif.org), parsed and verified using the ParseGBIF workflow (de Melo et al., 2024a, de Melo et al., 2024b). The resulting layer represents the difference (absolute value) between normalized GBIF/ParseGBIF and GIFT taxon richness. To remove excessively high taxon richness values produced by centroids within the GBIF/ParseGBIF richness map which were not removed by the ParseGBIF workflow, we capped it by the maximum GIFT taxon richness. We recognize that both data stes have limitations. GBIF occurrence data is highly biased and sparse, while GIFT models may be distorted due to the simplifications needed for global-scale data compounded by the biases in the underlying GBIF occurrence data identified above. However, when combined, these datasets provide the best available estimate of mismatch between sample effort and expected species diversity.

We considered including predicted β-diversity (species turnover) to prioritise areas (Socolar et al. 2016) where species-turnover is higher but ultimately excluded this variable because we could not find data layers in which we had confidence. For example, Moulatlet et al.’s maps of phylogenetic beta diversity (Moulatlet et al.2023) are counter-intuitive, indicating little decline in β-diversity with latitude until the boreal zone and proposing hotspots on the Baltic coast, and the Sahara Desert (Moulatlet et al., 2023, Fig. 1). In the absence of the available data we considered calculating beta diversity ourselves, using the ParseGBIF dataset (de Melo et al., 2024a) but felt that the result would be strongly biased by low and very uneven sampling density (Fig. 1).

To achieve a uniform map extent and grid cell size within each selected area, all variables were resampled to a resolution of 30 arc min (ca 50 km at the Equator) using bilinear interpolation and transformed to an equal area projection (Supplement 2.3). Within terrestrial areas, all “no data” values were replaced by zero, because TOPSIS cannot handle the former

### Tree-dominated vs grassland/deforested areas

Energy, water, and biogeochemistry patterns are different between forest and grassland (Breshears, 2006). Forest and grassland also differ in the ecosystem services they provide (Millennium Ecosystem Assessment, 2005). We prioritized areas within forest and grassland separately so as to avoid areas of forest and grassland being inadvertently discriminated against by the analysis. To separate the areas of predominantly grassland from forest we used the Global Land Cover-SHARE (GLC-SHARE) database created by FAO (https://data.apps.fao.org/). In each layer, pixel values represent the percentage of density coverage in each pixel of the land cover type. For the purposes of the project, we used two layers, “grassland’ and “tree covered area’, which correspond to areas with a high density of non-woody or woody plants respectively, with a cut-off of 10% basal area. As a result, the layers overlapped when there was a mixture of grassland and tree forest cover.

While this approach may misclassify some non-agricultural deforested areas as grassland, such instances are readily identifiable by consulting field experts and using mapping tools like Google Earth. Importantly, these deforested landscapes often correspond to former biodiversity hotspots that may harbour high proportions of threatened species or overlooked relict populations (Monro et al., 2001). The specific criteria for selecting local experts are detailed in Supplement 4

### Area prioritization

To prioritize areas for new data acquisition, we used a multi-criteria-based decision-making Technique for Order Preference by Similarity to Ideal Solution (TOPSIS; Hwang Ching-Lai & Yoon, 1981) implemented in the R package “topsis”. TOPSIS chooses the best alternative which has the shortest Euclidean distance from the positive ideal solution and the greatest, from the negative ideal solution (Table S2.1). TOPSIS has been used successfully in a variety of fields, including environmental sciences (Mitra et al., 2023). The advantages of TOPSlS are its simplicity, transparency, and adaptability.

When assigning weights to the selected variables, we were informed by our assessment of variable importance based on the ecological and macroecological literature. For example, terrain roughness as a driver of species-richness was assigned a positive weight based on Stein et al. (2014) and this was set at 0.2 following a discussion with the authors of the relative importance of the different data layers. We used an analytic hierarchy process with pairwise comparison (Teknomo, 2012). The importance of all factors was compared on the scale 1 to 10. The resulting comparison matrix (Table S2.2) was processed using the R package “SpatMCDA” (Wang et al., 2024) to obtain variable weights (Table S2.3). Wang et al. (Wang et al., 2024) demonstrated that the change in variable weights in the range -20% to 20% was of minor importance to the model outcomes, therefore we did not explore variability in weights.

We applied TOPSIS at one global and five regional scales (Table S2.4) and projected the scores onto regional maps with ca 50×50 km grid cells. The grid cells prioritized by TOPSIS for further sampling efforts had values close to 1, while the cells with low priority had values close to zero. Because the spread of TOPSIS scores varied across regions and vegetation types (grassland or forest), when visualising TOPSIS scores we used an equal area method so that the areas spanned by each colour occupied the same extent of the map.

### Field expert selection and assessment

The priority areas identified by the TOPSIS algorithm were reviewed by plant scientists with specific regional or national field expertise. Twenty-seven experts were selected, covering 19 countries or subregions. Selection was based on their scholarly publications and contributions to biological collections pertinent to their respective regions of expertise. Feedback took the form of responses to specific questions. To validate the model’s output, a panel of twenty-seven botanical experts with specific regional expertise (covering 19 countries and subregions) was consulted. Each expert was provided with the area ranking maps (Supplement 4, Fig. 3–8), the complete methodology, and maps of the underlying environmental variables (Supplement 1). They were asked to: **(i)** assess the value of the identified high-priority areas for guiding future sampling efforts; **(ii)** support or contest the overall prioritization for their region of expertise; and **(iii)** delineate any specific subareas where their assessment differed. The extent of agreement in feedback was calculated as a Consensus Proportion (S3) as opposed to more formal measures such as applying a formal inter-rater reliability statistic like Cohen’s Kappa or Fleiss’ Kappa. This was because those metrics are best suited for situations where the areas reviewed are discrete, predefined categories. Whilst this was he case for our prioritizated areas, it is not for the disputed areas which will be sub- and so, smaller, areas. The Consensus Proportion equals the number of supported areas divided by sum of the number of supported and unsupported areas. Following expert feedback, we refined our methodology by stratifying the dataset into tree-dominated and non-tree-dominated areas, which guided a revised prioritization analysis.

**Fig. 3.**
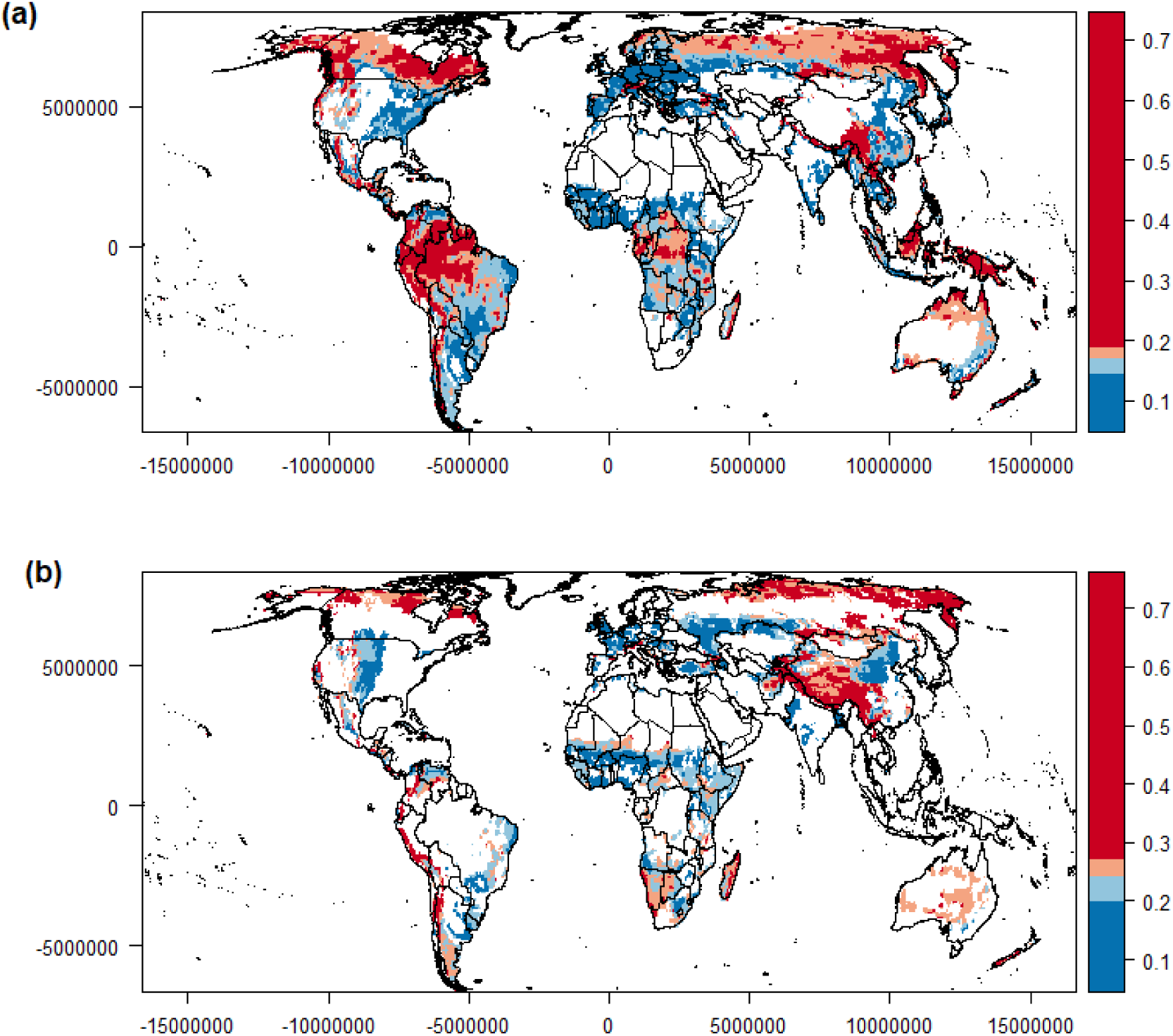
Area ranking for future sampling efforts, global extent: (a) tree-dominated areas; (b) non-tree-dominated areas. Red represents the areas of highest priority for data acquisition; dark blue, the areas of lowest priority; light blue and orange represent the areas of medium priority; white represents areas with no data. Each priority rank (colour) occupies ca 25% of the total area of either grassland or forest cover. Map projection is Eckert IV (equal area). Map units are meters. Map created with custom R script. Base map source: ESRI (http://www.esri.com/data/basemaps, © Esri, DeLorme Publishing Company).

**Fig. 4.**
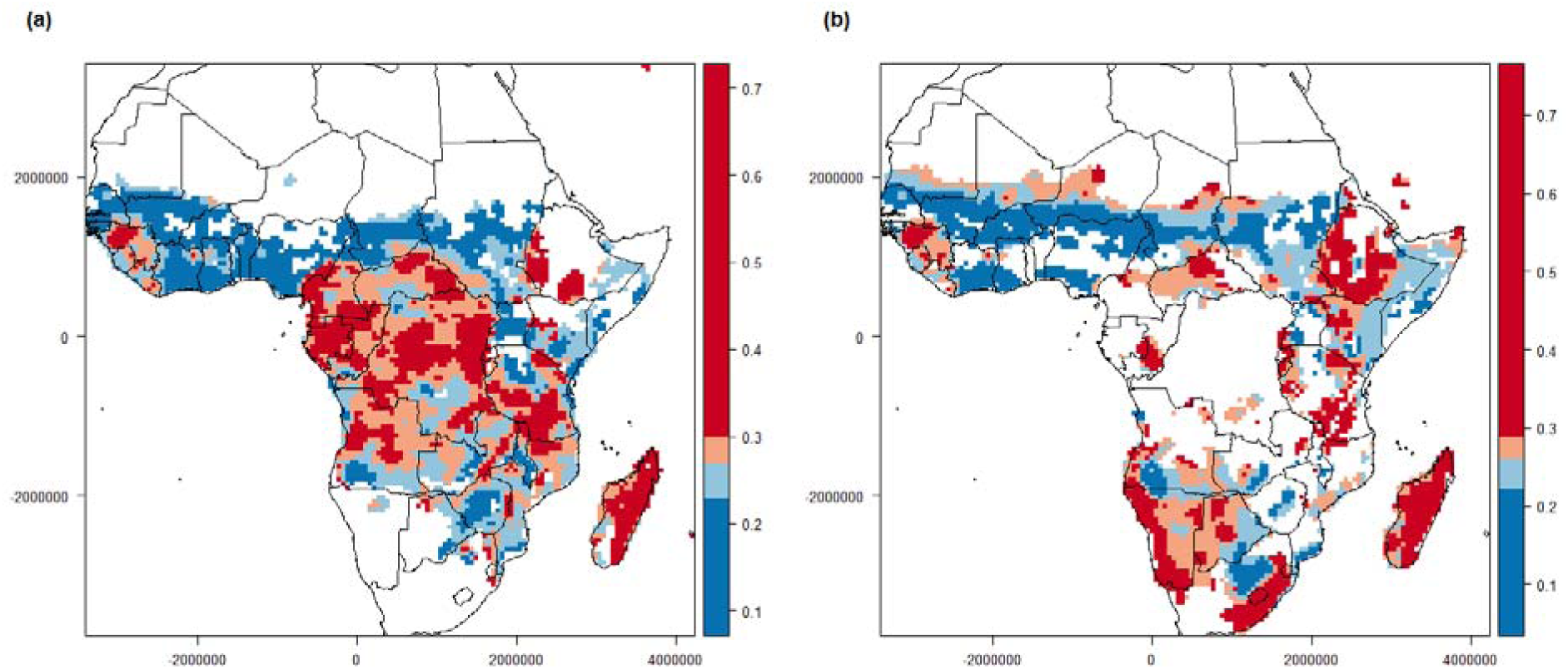
Area ranking for future sampling efforts, Africa and Madagascar: (a) tree-dominated areas; (b) grassland/deforested areas. Red represents the areas of highest priority for data acquisition; dark blue, the areas of lowest priority; light blue and orange represent the areas of medium priority; white represents areas with no data. Each priority rank (colour) occupies ca 25% of the total area of either grassland or forest cover. Map projection is Equatorial Lambert azimuthal (equal area). Map units are meters. Map created with custom R script. Base map source: ESRI (http://www.esri.com/data/basemaps, © Esri, DeLorme Publishing Company).

**Fig. 5.**
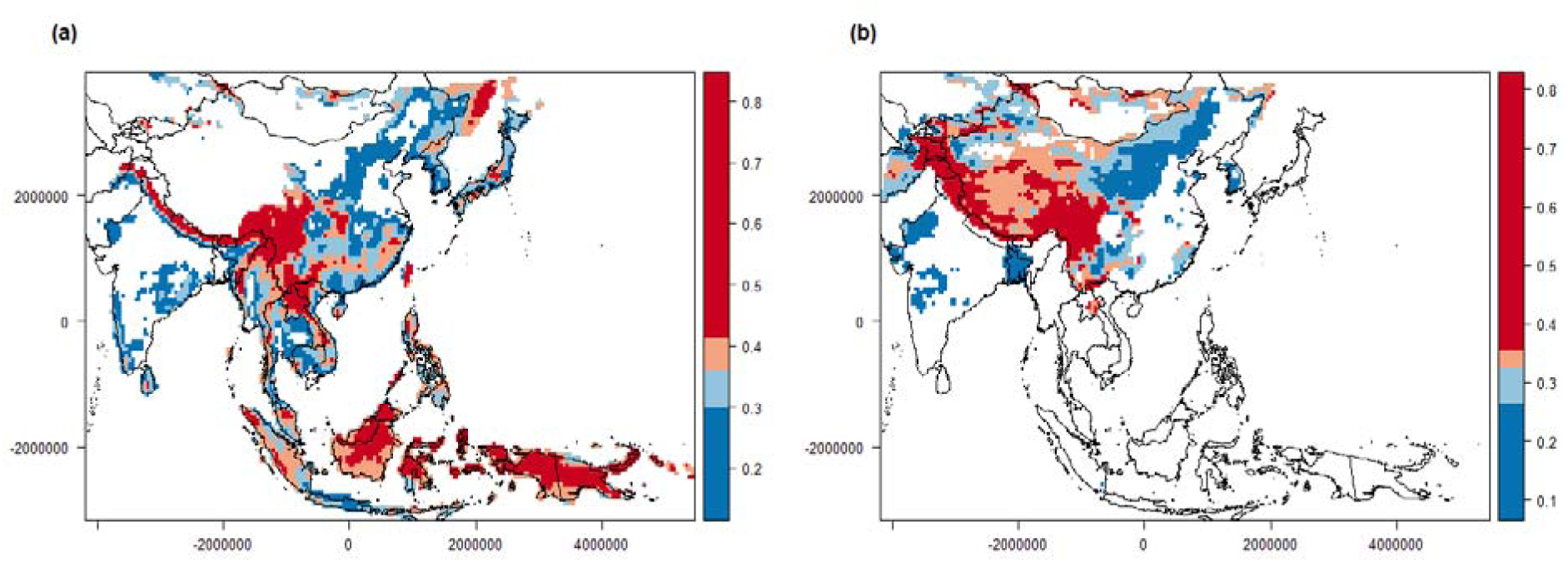
Area ranking for future sampling efforts, East, South and Southeast Asia: (a) tree-dominated areas; (b) grassland/deforested areas. Red represents the areas of highest priority for data acquisition; dark blue, the areas of lowest priority; light blue and orange represent the areas of medium priority; white represents areas with no data. Each priority rank (colour) occupies ca 25% of the total area of either grassland or forest cover. Map projection is Albers conic (equal area). Map units are meters. Map created with custom R script. Base map source: ESRI (http://www.esri.com/data/basemaps, © Esri, DeLorme Publishing Company).

**Fig. 6.**
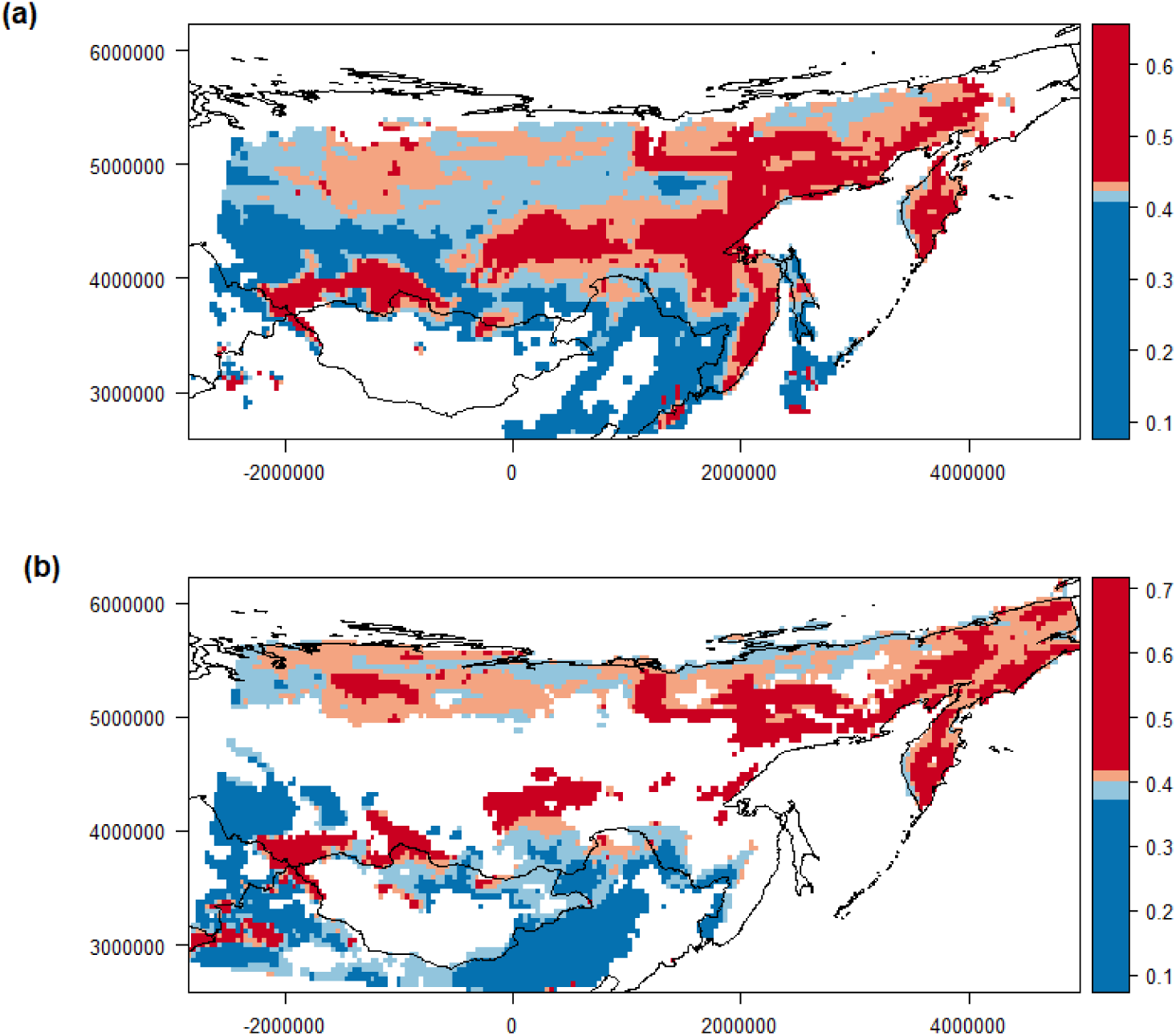
Area ranking for future sampling efforts, Siberia and Russian Far East: (a) tree-dominated areas; (b) grassland/deforested areas. Red represents the areas of highest priority for data acquisition; dark blue, the areas of lowest priority; light blue and orange represent the areas of medium priority; white represents areas with no data. Each priority rank (colour) occupies ca 25% of the total area of either grassland or forest cover. Map projection is Albers conic (equal area). Map units are meters. Map created with custom R script. Base map source: ESRI (http://www.esri.com/data/basemaps, © Esri, DeLorme Publishing Company).

**Fig. 7.**
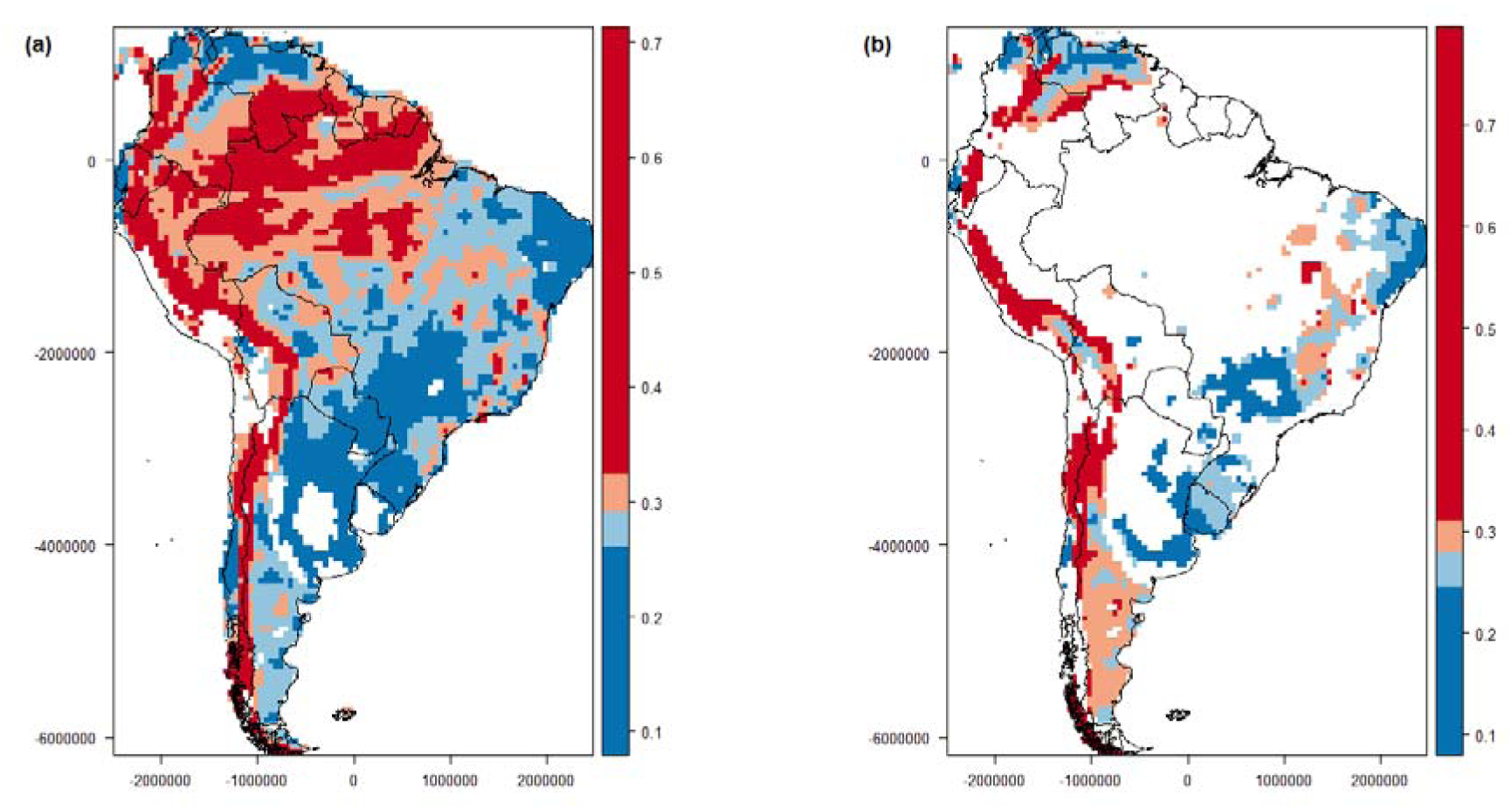
Area ranking for future sampling efforts, South America: (a) tree-dominated areas; (b) grassland/deforested areas. Red represents the areas of highest priority for data acquisition; dark blue, the areas of lowest priority; light blue and orange represent the areas of medium priority; white represents areas with no data. Each priority rank (colour) occupies ca 25% of the total area of either grassland or forest cover. Map projection is Eckert IV (equal area). Map units are meters. Map created with custom R script. Base map source: ESRI (http://www.esri.com/data/basemaps, © Esri, DeLorme Publishing Company).

**Fig. 8.**
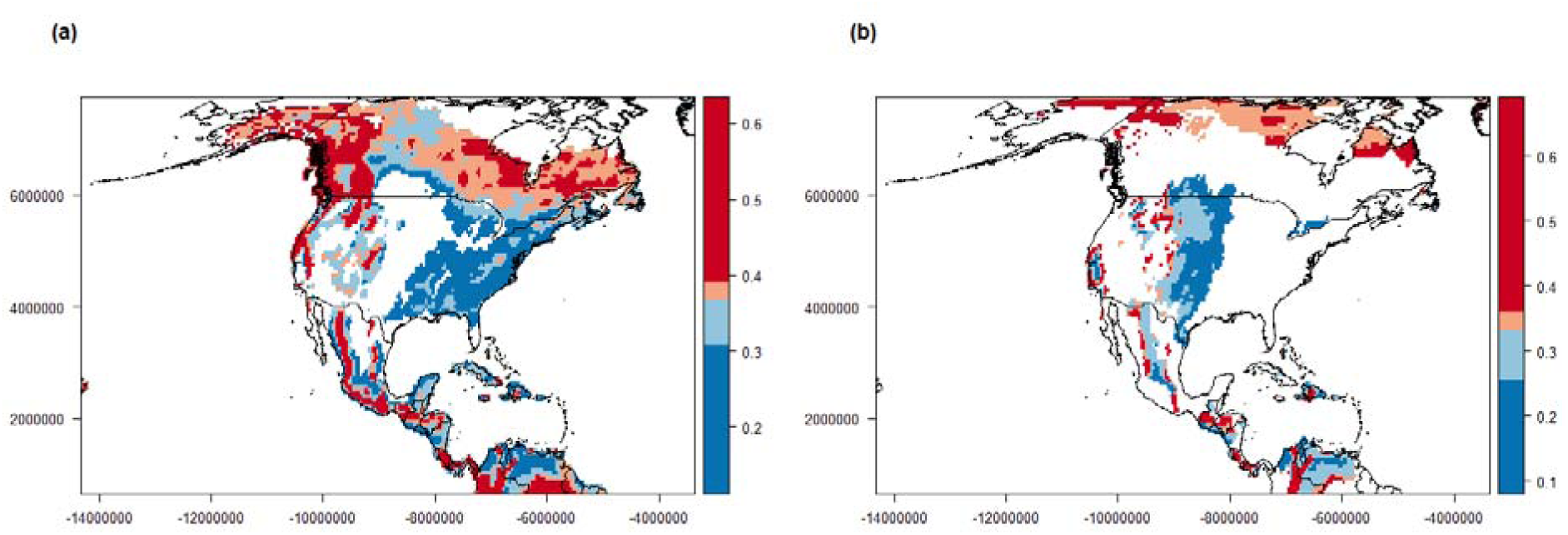
Area ranking for future sampling efforts, North America, Central America and the Antilles: (a) tree-dominated areas; (b) grassland/deforested areas. Red represents the areas of highest priority for data acquisition; dark blue, the areas of lowest priority; light blue and orange represent the areas of medium priority; white represents areas with no data. Each priority rank (colour) occupies ca 25% of the total area of either grassland or forest cover. Map projection is Eckert IV (equal area). Map units are meters. Map created with custom R script. Base map source: ESRI (http://www.esri.com/data/basemaps, © Esri, DeLorme Publishing Company).

## Results

We present the results for the global extent (Fig. 3) and five regional spatial extents: Africa and Madagascar, East Asia, Siberia and the Russian Far East, South America, and North America, Central America and the Antilles (Figures 4 – 8).

For all analyses, correlations between environmental variable did not exceed 0.7, except in the Siberia and Russian Far East and North and Central Americas, where the variables “Global human modification of terrestrial systems” and “Sampling bias” were negatively correlated at 0.81 and 0.75, respectively; TOPSIS scores were below 0.85 (Supplement 2).

Supplement 3 provides a summary of the field expert reviewer feedback, highlighting disagreement with the results of our TOPSIS analysis. The proportion of consensus showed support ranging from 85% to 100% for the prioritization of regions, with East, South and Southeast Asia having the greatest disagreement (see S3, ‘Summary of Proportion of Consensus’

### Global extent

In tree-dominated areas (Fig. 3a), global analyses identified the highest priority areas as: the Andes, Amazonia, and Atlantic Forest subregions in South America; parts of Mesoamerica (including Costa Rica, Panama, and Guatemala); various regions of North America (including the Sierra Madre, US-Canada Pacific coast forests, and parts of the Taiga and Hudson Plain); Central Africa (Congolian region), parts of East (Zambezian region) and southern Africa (Southern African region), and Madagascar; the Himalayas, parts of Southeast Asia and Australasia (including Borneo, Sulawesi, New Guinea, and parts of Australia and New Zealand); and several regions of Eurasia (including the Dolomites, Caucasus, Siberia, Altai, and Kamchatka).

In non-tree-dominated areas (Fig. 3b), global analyses prioritized the following regions: South America (Andes, Guyana Shield); Mesoamerica (Costa Rica, Guatemala); North America (Sierra Madre, US-Canada Pacific coast forests, Taiga); Africa (Congolian, Zambezian, Southern African regions, including Central African Republic, Namibia, Tanzania/Mozambique, Botswana, Madagascar); Asia/Australasia (Himalayas, Tibetan Plateau, Laos, island of Taiwan, Australia, New Zealand); and Eurasia (Iberia, Dolomites, Caucasus, Siberia, Altai, Kamchatka).

### Africa and Madagascar

Field expert feedback largely supported 95% of the highest prioritized areas (S3).

Our analyses of tree-dominated areas (Fig. 4a) prioritized future data acquisition in several African biogeographic regions (Congolian, Zambezian, Djibouti, and Somalian; Linder et al., 2012). These prioritized areas include parts of Liberia, Guinea, Cameroon, Central African Republic, Democratic Republic of Congo, Gabon, Republic of the Congo, Angola, Tanzania, northern Mozambique, Ethiopia, Sudan, and Kenya, as well as the humid forests of Madagascar. Field expert identified the lack of high-priority areas in Liberia, and doubts about the ranking for Ethiopia raised (S3).

Our analyses of non-forest dominated areas (Fig. 4b) prioritized future data acquisition in several in several African biogeographic regions (*sensu* Linder et al., 2012; Sudanian, Congolian, Zambezian, Djibouti, Somalian and South African). These prioritized areas include the Fouta Djalon (Guinea), the grasslands of Ethiopia, parts of Kenya, Somalia, and Tanzania, the Atlantic coast of Namibia, the Eastern Cape and KwaZulu-Natal of South Africa and Lesotho, and Madagascar. Reviewer feedback largely supported the top 25% of prioritized areas, but the lack of prioritization for coastal grasslands/thickets in coastal Congo-Brazzaville, Gabon, DRC, Sierra Leone, and Liberia was noted (S3). Field expert reviewers disagreed with the classification for part of the Ethiopian region (sensu Linder et al., 2012), concluding that our analyses had underprioritized Acacia-Commiphora forests in the southeast of the country.

### East, South and Southeast Asia

Field expert feedback supported 85% of the highest prioritized areas (S3).

Within the tree-dominated extent of East, South and Southeast Asia (Fig. 5a), the major prioritized regions for additional data acquisition were the Himalaya mountains, the Tibetan plateau, the Guizhou – Yunnan plateau, subareas of the Sunda Shelf, Wallacea and all of the Sahul Shelf including parts of Western Laos, Northern Malaysia, Aceh (Indonesia), western Sumatra (Indonesia), Central and Northern Borneo, Sulawesi (Indonesia), the Banda Arc (Indonesia), North Maluku (Indonesia) and New Guinea. Field expert reviewers suggested that Timor was under-prioritised; that in Borneo the southern extent should be a higher priority than the northern extent; and that in New Guinea the southern Savannahs and Mount Jaya should not be the highest priority (S3).

For non-tree-dominated areas in East, South and Southeast Asia (Fig. 5b), the major prioritized regions for additional data acquisition were the Tian Shan, Hindu Kush, Karakorum, and Himalayan mountains and the Tibetan Plateau including parts of Afghanistan, Kyrgyzstan, Uzbekistan, Pakistan, India, Mongolia, China, and Laos.

### Siberia and the Russian Far East

Field expert feedback supported all of the highest prioritized areas (S3).

Within the forest-dominated extent the highest priority areas were the Ural Mountains (including the territory of the Sakha Republic, Chukotka Autonomous Region, Magadan Region) and Kamchatka, the Altai and Tuva Republics, and Buryatia (Fig. 6a), with an addition of the Sakhalin Region for the non-forest dominated extent (Fig. 6b).

### South America

Field expert feedback supported 92% of the highest prioritized areas (S3).

For tree-dominated areas our analyses (Fig. 7a) identified the Andean forests and the Amazon basin, along with patches of inland Atlantic rain forest, Caatinga, and Cerrado, as the highest priority areas for future data acquisition.

For non-forest dominated areas our analyses (Fig. 7b) prioritized high elevation grasslands of the Andes and subareas of the Cerrado for future data acquisition. Field expert reviewers queried the absence of the Pampas grasslands adjacent to the Uruguay-Brazil border, as well as the Pampas in the Beni and the Chiquitania in Bolivia, as high priorities.

### North America, Central America and the Antilles

Field expert reviews were undertaken for Mesoamerica only and supported all of the highest prioritized areas (S3)

For tree-dominated areas our analyses (Fig. 8a) at the regional scale recovered the marine west coast forests, the full extent of the north west forested mountains and subareas of the northern forests of North America, western and eastern subareas of taiga, the Hudson Plain, subareas of the northern American desert in Mexico, the tropical dry forests of Mexico, the seasonal dry forests of northeastern Cuba and southwestern Dominican Republic, the montane forests of Mesoamerica, and the tropical forests of Costa Rica and Panama as the highest priority areas in North America, Mesoamerica and the Antilles. Field expert reviewers queried the absence, as a high priority, of the coastal Pacific forests of El Salvador and Guatemala (S3).

For non-tree-dominated areas (Fig. 8b) our analyses at the regional scale recovered the western and eastern subareas of taiga, the Arctic Cordillera, the Great Plains, subareas of the northern American desert in Mexico, Mediterranean vegetation of Mexico, subareas of dry forest in southern Mexico, southwestern Dominican Republic, highlands of Mesoamerica, as the highest priority areas in North America, Mesoamerica and the Antilles.

## Discussion

### A validated and scalable framework for identifying priority areas

The multi-criteria decision method TOPSIS is a transparent, computationally efficient, and highly adaptable framework for prioritization (Taherdoost & Madanchian, 2023). Its versatility, the ability to be applied to any taxon or combination of taxa, makes it particularly suitable for developing the national-level prioritizations required by the Global Biodiversity Framework (GBF). A key strength of this approach is its capacity to translate expert opinion into explicit and quantitative criteria, which in turn facilitates clear communication with policymakers and funders. Using defined variables, adjustable weighting, and data compartmentalization, the method produces prioritized maps and lists that can be tailored to specific national or stakeholder needs. Furthermore, this transparency ensures that if a result is questioned, the underlying data and weighting can be interrogated, adjusted, and the analysis rerun. This inherent traceability offers a distinct advantage over AI-based ‘black box’ approaches, where anomalous results can be difficult to interpret and computationally expensive to correct.

Regional analyses were compared to global analyses as a preliminary assessment of scalability. The largely congruent global and regional results support the utility of our approach by increasing confidence in the identification of key priority areas, ensuring their significance is reflected at both broad and local levels. An exception was Angola, which is identified more clearly as a priority in the regional analysis, compared to the global one. We also found substantial overlap between the highest priority areas identified for both forest-dominated and non-forest-dominated regions, with the latter consistently appearing as a subset of the former.

Our regional analyses, validated by field expert review, demonstrated strong agreement between TOPSIS-derived priority areas and local field knowledge (S3). Where disagreement did occur, this was mostly in relation to the non-forest-dominated map, and specifically to their conflation with deforested areas. Whilst deforested areas are widely considered a low-priority for conservation, and by extension, data acquisition, deforested areas can harbour threatened relict populations of native flora. For example, according to Monro et al. (2001), approximately one-third of the original tree flora persist in shade coffee farms in El Salvador, highlighting the potential ecological significance of such altered landscapes.

In relation to forest-dominated areas, the area of highest priority for Borneo (Fig. 5a) and Ethiopia were queried (S3). One of the field experts suggested that the highest priority areas for Borneo should be in the more human-impacted south of the island, rather than in the north as identified by our analysis. A review of the variables used (Fig. S1) suggests that our result may have been influenced by the greater frequency of threatened species (Fig. S1.12) and terrain roughness (Fig. S1.6) in the north of Borneo. The advantage of the TOPSIS approach is that the weighting of these layers can be adjusted. In the case of Ethiopia, reviewers disagreed with the Ogadden (in the Somali Region of Ethiopia and Somalia, Fig. 4a) not being ranked as a high priority, given that *Acacia*-*Commiphora* woodland, is poorly known and rich in endemic species (S3). A review of the variable layers indicates that most of this area was classified as non-forest dominated and subject to relatively high sample bias (Fig. S1.13), low human modification and with a close match between observed and predicted species-richness (Fig.S1.15). In a more targeted analysis, the above could be mitigated by increasing the weighting applied to sample bias and lowering the weighting applied to the mismatch between observed and expected diversity (Fig.S1.15).

Whilst our aim is to prioritise areas for future data acquisition (exploration, digitisation and the curation of existing data), there was general concordance between areas ranked within the highest 25% priority for future data acquisition, and those identified by previous studies as hotspots for conservation (Myers et al., 2000), darkspots for future exploration (Ondo et al., 2024) or centres of vascular plant species richness (Cai et al., 2023; Sabatini et al., 2022). Notably, our analysis identifies several high-priority regions—including Amazonia, and parts of Central and Eastern Africa—that are often overlooked in other global conservation assessments. Amazonia provides an obvious example. Despite being one of the world’s most species-rich and threatened forested areas, it is consistently missed in global prioritizations (ter Stege et al., 2006; Vera et al., 2023). We attribute this to methodological differences. Many previous analyses rely on raw species occurrence data, where severe under-sampling and geographic biases can distort model outputs. In contrast, our approach uses occurrence data primarily to model sampling bias and the mismatch between observed and expected diversity, rather than as a direct measure of distribution. This likely explains why our results, alongside those of Sabatini et al. (2022) who used plot data to identify well-sampled proxy variables, successfully highlight Amazonia’s importance. Other discrepancies can be attributed to different prioritization criteria; for instance, Myers et al. (2000) likely did not classify Amazonia as a hotspot because their framework placed greater emphasis on immediate threats from human modification.

A similar situation is evident for northern boreal biomes, where we identify the Putorana Plateau and Canada’s Hudson Plain as high priorities for data acquisition. This assessment aligns with that of Sabatini et al. (2022), who classified these same regions as “data-deficient.” Our prioritization of areas for additional occurrence data acquisition, both in tree-dominated and non-tree-dominated regions, strongly agreed with the findings of Sabatini et al. (Sabatini et al., 2022). For tree-dominated areas, our priorities aligned with regions above the 95th percentile for vascular plant species richness (Sabatini et al., 2022. Figs 1), while for non-tree-dominated areas, they matched regions above the 95th percentile for species richness or those identified as data-poor (Sabatini et al., 2022. Figs 2). This congruence highlights the significant influence of area species-richness on our prioritization strategy, with more species-rich areas, as identified by Sabatini et al. (2022) receiving higher priority.

Our results differ from the identification of darkspots as global collection priorities for the discovery of new species acquisition of additional occurrence data (Ondo et al., 2024). As above, we identified the Amazon as amongst the highest priority areas for filling knowledge-gaps in forest-dominated areas of South America (Fig. 7a). The Amazon was not, however, classed as a high priority for future collecting by Ondo et al. (Ondo et al., 2024). In addition, Ondo et al. (Ondo et al., 2024) identified South Africa, regarded as one of the best documented floras (the Cape Flora; Cowling et al., 2009) as the only African darkspot (Ondo et al., 2024), whilst in contrast, we identified subareas of tropical moist and dry vegetation across Western, Central and eastern Africa (Figs 3a, b) as most in need of additional sample effort.

### Limitations of the study

While the compartmentalization of data layers and prioritization areas is essential for effective prioritization, classifying areas as strictly forest-dominated or non-forest-dominated presents inherent challenges. We addressed this by adopting the FAO’s Global Land Cover-SHARE (GLC-SHARE) database (https://data.apps.fao.org/). However, the definition of “forest-dominated” includes an arbitrary element, and deforested areas are classified as non-forest. Furthermore, aggregation to a 50 x 50 km cell size resulted in some areas with forest cover being classified as non-forest and vice versa. These limitations, however, only apply where the maps are used independently of each other, whilst our intention is that both maps are used in-tandem, priority areas from each map being overlayed on a single map.

The 250 km² cell size employed here likely resulted in the aggregation and subsequent loss of spatially restricted vegetation types, such as those found in littoral or riparian zones. This limitation is exemplified by the analysis’s failure to resolve white sand coastal vegetation in Ghana, Sierra Leone, and Liberia, which, due to its narrow spatial extent, was significantly smaller than the cell resolution (see S3). This can be overcome in smaller-scale, e.g. national prioritizations, through the application of a smaller cell-size.

Including β-diversity (Socolar et al., 2016) or complementarity (Bush et al., 2016) in the TOPSIS analysis would improve the prioritization of areas for biodiversity documentation. However, obtaining high-resolution β-diversity estimates at regional, or the national scale—the level for implementing Global Biodiversity Framework (GBF) actions—is often challenging given the quality of existing occurrence data for plants (Fig.1).

Lastly, the simplicity of our method over more complex models brings advantages in terms of adjustment, interpretability, and communication. However, they may not adequately capture the complexity of the real world, such as changes in land use and climatic stresses over time that may alter the threat level to species in particular regions, for which machine learning methods can demonstrably outperform simpler models and human intuition (<u>Silvestro et al., 2022</u>). Despite its simplicity, however, our method still carries a relatively large number of parameters and weightings, chosen based on our informed choices but subject to further exploration and sensitivity analyses.

### Strategies for acquiring additional occurrence data

Whilst documenting the vascular plant diversity of the world’s biodiversity rich regions with primary data by 2030 is likely untenable, obtaining the minimum number of point occurrences to generate effective SDMs for all species should be tractable through a campaign of data generation that targets sample biases, threatened areas and the drivers of species richness.

There are four main routes for generating this data, 1) targeted on the ground inventories; 2) the integration of the extensive network of vegetation plots and preserved specimen collections (many of which are not yet digitally available), both with respect to species identifications, and the documentation of the non-tree flora; 3) the harvesting of occurrence records currently unusable because they lack either spatial coordinates (*ca* 32 million collection events in GBIF) or a species identification (*ca* 6 million collection events in GBIF); and 4) the digitisation of specimens in hitherto overlooked collections (e.g. BUD, LEN, local herbaria). While approaches 3) and 4) can increase sampling efforts, they are unlikely to mitigate the sample biases present in GBIF, as they reflect the same historical processes. Only approaches 1) and 2) have the potential to increase sample effort while addressing sampling bias.

An additional avenue to address the challenges in generating high-quality occurrence data lies in fostering interdisciplinary collaboration and leveraging emerging technologies. Integrating remote sensing, artificial intelligence, and machine learning with traditional field-based methods could enhance the accuracy and efficiency of data collection. For instance, high-resolution satellite imagery has been used to identify and monitor biodiversity hotspots, providing large-scale environmental data that complement ground inventories (Pettorelli et al., 2014; Turner et al., 2003). AI-driven species identification algorithms and deep learning techniques offer powerful tools for automating the identification of species and analyzing ecological data (Christin et al., 2019).

Generating additional verifiable occurrence data *de novo* (source 1) will require significant improvements in species identification, the accessioning process, and digitisation to meet the 2030 target. Current developments in bioinformatics, robotics and machine learning provide for occurrence data generation and migration into a portal instantaneously from the field, as suggested by the Xprize Rainforest Names Biodiversity Tech Competition (2024). Identification to species and verification would, however, require that associated workflows (permits, shipping/ unpacking, registration, identification, mounting, digitisation), be accelerated considerably.

The integration of vegetation plot networks and preserved specimen collections (source 2), offers great potential as many of the verifiable occurrence data exist as vouchers in the local institutions tasked with maintaining them, and the expansion of the species sampled to include herbs, shrubs, treelets, and epiphytes would establish datasets of greater predictive value (Baker et al., 2017). Vegetation plot data (or “grass-root data”) provide crucial information on species co-occurrence, invaluable for understanding plant community composition, distribution, structure, function, and evolution. However, barriers such as erroneous or missing geolocations and inconsistent plant names hinder their use (Wiser, 2016). These challenges can be mitigated through voucher integration, greater collaboration with taxonomists and botanical institutions, and the use of named entity recognition in multi-lingual models (Fetahu et al., 2022).

The harvesting of spatial data for occurrence records which lack coordinates, but which have precise text-based locality information (source 3), would increase the number of useable occurrence records by at least one third, relatively quickly and economically. Whilst vascular plant occurrence records are increasingly aggregated over time, Feeley & Silman (Feeley & Silman, 2011) conclude that additional sites are nonetheless added, making it likely that harvesting spatial data would lead to improved sample coverage (Smith et al., 2023). The targeted digitisation of subnational herbaria (source 4) with coverage for the highest priority areas, would also represent an approach to acquiring additional occurrence data, congruent with the 2030 deadline for the CBF targets. However, both approaches are subject to the same biases and intrinsic limitations inherent in historical sampling efforts (Delves et al. 2023., Di Cecco et al., 2021; Geurts et al., 2023).

Our research aims to support the generation of a plant occurrence dataset of sufficient quality to underpin the implementation of the GBF. Specifically, we aimed to facilitate reduction in sample bias, stimulate increased sample effort in areas that best target knowledge gaps and encourage institutions to coordinate their efforts. By doing so in an analytical context that factors in threats to areas and ecosystem services, we aspire to offer funders a clearer understanding of the potential impact of proposed inventories, either plot- or expedition-based and so increase their motivation to support them.

Because additional primary data is needed (Antonelli et al., 2023; Antonelli et al., 2024) the botanical community must coordinate its efforts, to invest significantly in the generation of new data (source 1), or to support the integration of plot vouchers (source 2), both of which will be necessary to achieve a spatial occurrence data set for vascular plants that can support the GBF. One example of where such collaboration and coordination are taking place is the World Flora Online consortium (WFO 2024), a collaboration between 52 botanical institutions. Whilst WFO is currently focussed on the dissemination of taxonomic knowledge, it could be envisaged that, as in the WFO Taxonomic Expert Networks (TENs), geographical equivalents could be established for the generation and curation of occurrence data, potentially in collaboration with GBIF. The WFO could serve as a focus for establishing strategic collaborations between institutions to achieve this.

Funding for data acquisition, currently very scarce, even in the economically powerful regions, would also need to increase considerably. Aubin et al. (Aubin et al., 2020) stress the importance of collaborative research and outline the major requirements for the effective data acquisition: “(a) a data management plan; (b) data archiving and description standards; (c) dedicated resources to project coordination and data provider support; (d) a clear objective/mandate; and (e) an explicit link to global data initiatives”. We propose that a prioritization framework such as presented here, would not only support the above but also provide a convincing case of the needs and benefits of funding such data acquisition.

By combining these innovations with traditional methods, botanical institutions can address data gaps more effectively and achieve the spatial coverage necessary to implement the GBF. Such technological advancements, alongside collaborative efforts, have been recognized as critical for conservation and biodiversity research (Pimm et al., 2015).

## Data accessibility statement

The data that support the findings of this study are openly available from the data sources cited in the article. Parsed GBIF global plant occurrence data (unique collection localities) are openly available in figshare at 10.6084/m9.figshare.30264967.

## Supporting Information

Supplement 1. Environmental variables.

Supplement 2. TOPSIS model parameters and scores. Supplement 3. Field expert assessment.

Supplement 4. List of local vegetation experts.

Supplement 5. Technique for Order of Preference by Similarity to Ideal Solution (TOPSIS)

Unique collection localities of parsed GBIF global plant occurrence data are openly available in figshare at https://figshare.com/s/6e2d149636034c120370

