## Supplement 1 for "Setting priorities for the acquisition of primary plant occurrence data"

Supplement 1. Environmental variables.

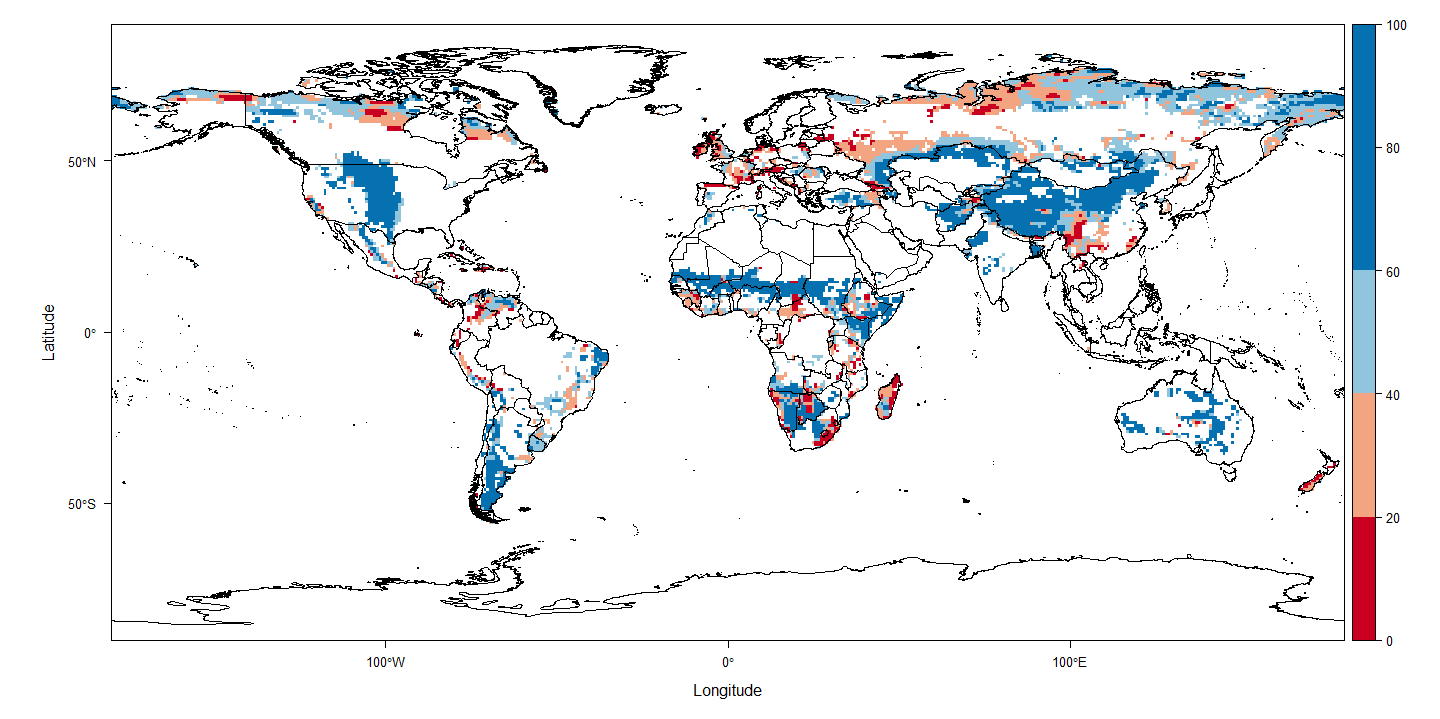

Fig. S1.1. Areas of global importance for conserving terrestrial biodiversity, carbon, and water: grassland. Base map source: ESRI (http:// www. esri. com/ data/ base maps, © Esri, DeLorme Publishing Company).

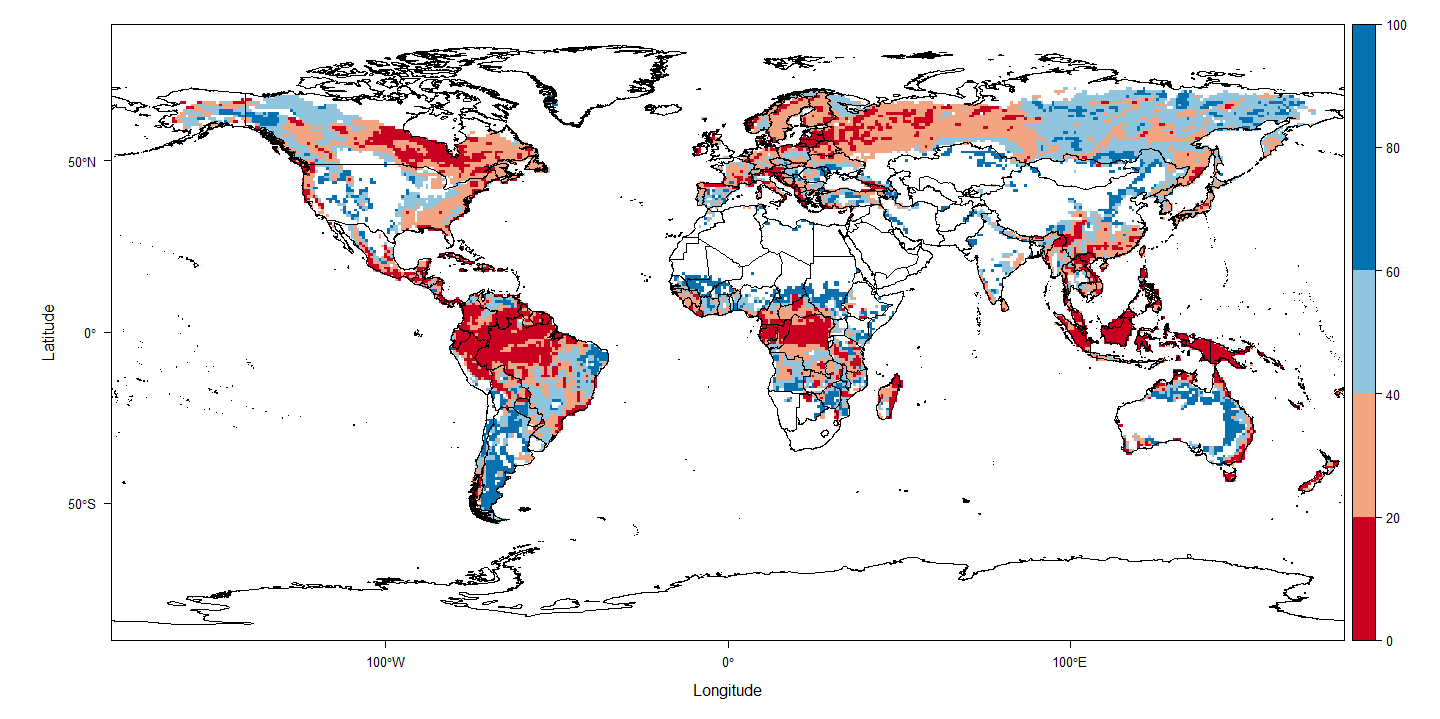

Fig. S1.2. Areas of global importance for conserving terrestrial biodiversity, carbon, and water: forest. Base map source: ESRI (http:// www. esri. com/ data/ base maps, © Esri, DeLorme Publishing Company).

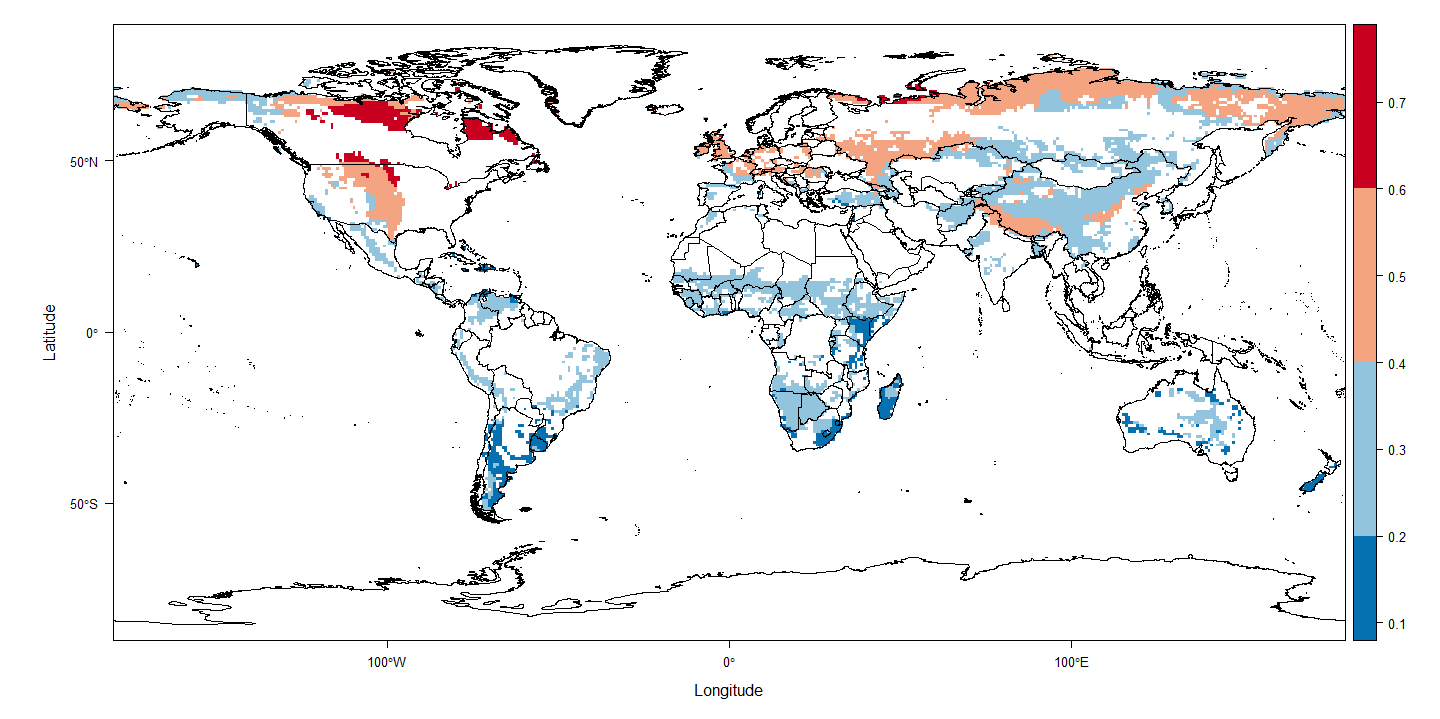

Fig. S1.3. Climate stability from Pliocene (3.3 Ma) to the present: grassland. Base map source: ESRI (http:// www. esri. com/ data/ base maps, © Esri, DeLorme Publishing Company).

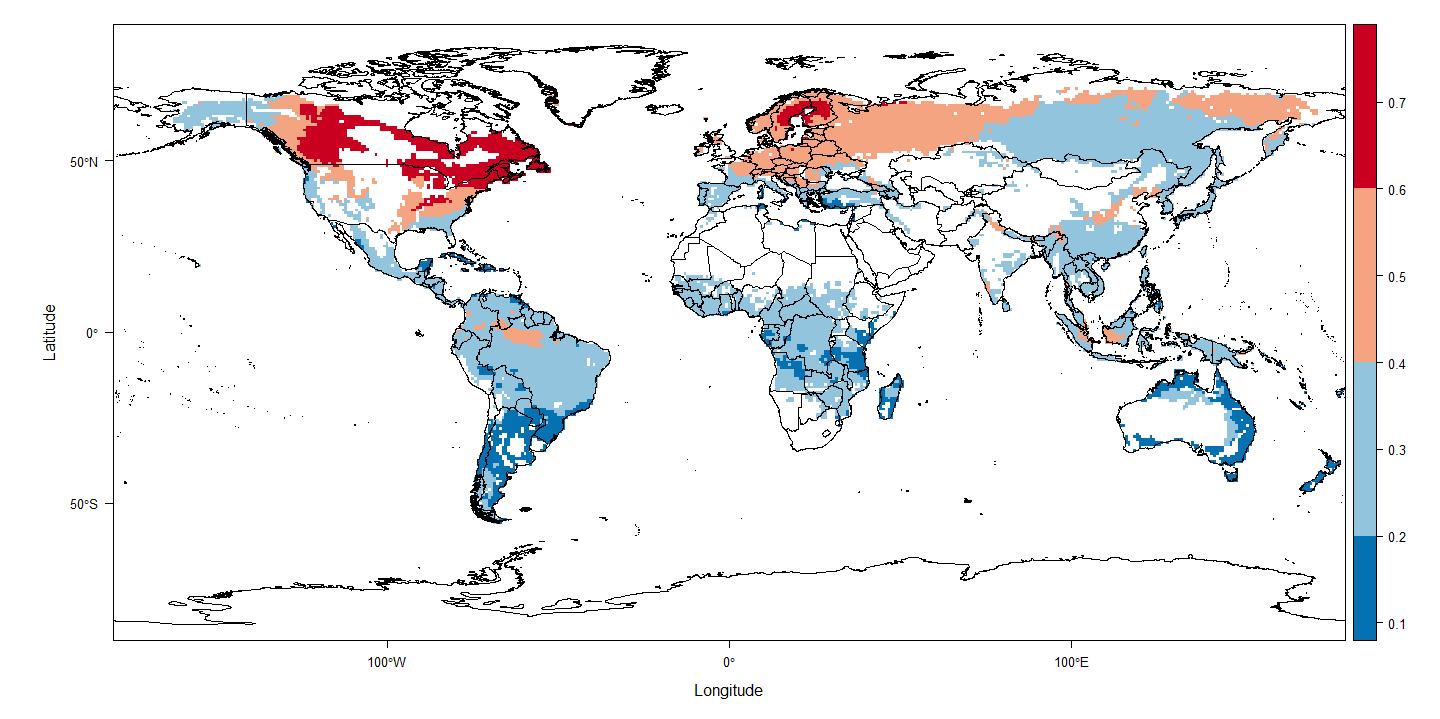

Fig. S1.4. Climate stability from Pliocene (3.3 Ma) to the present: forest. Base map source: ESRI (http:// www. esri. com/ data/ base maps, © Esri, DeLorme Publishing Company).

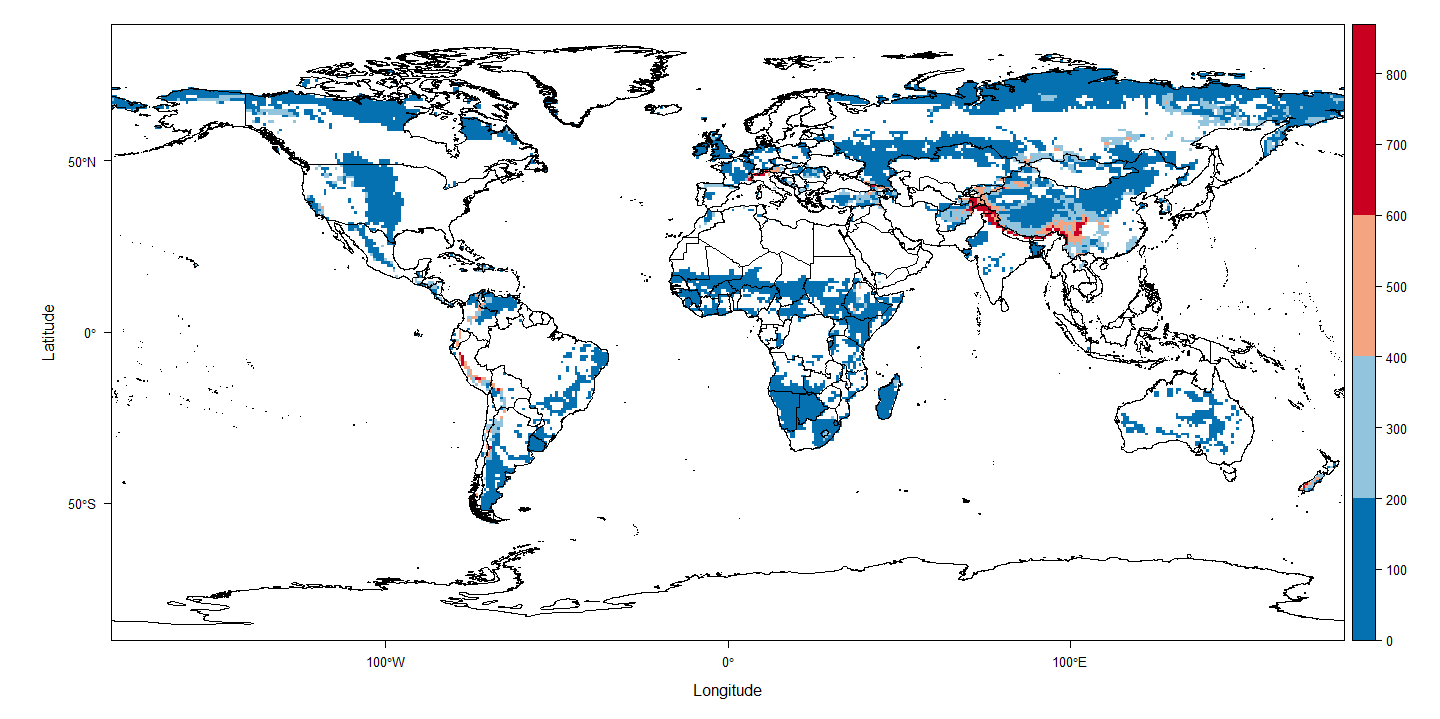

Fig. S1.5. Terrain roughness: grassland. Base map source: ESRI (http:// www. esri. com/ data/ base maps, © Esri, DeLorme Publishing Company).

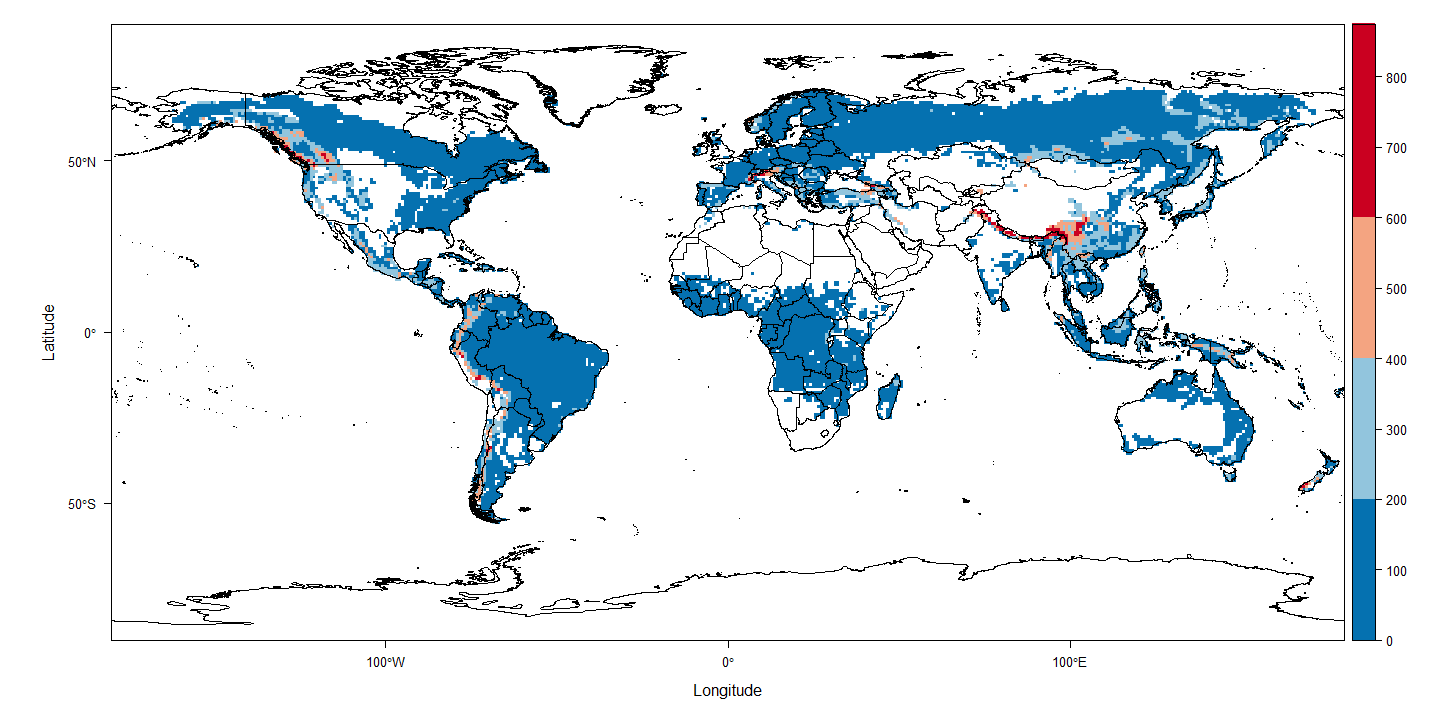

Fig. S1.6. Terrain roughness: forest. Base map source: ESRI (http:// www. esri. com/ data/ base maps, © Esri, DeLorme Publishing Company).

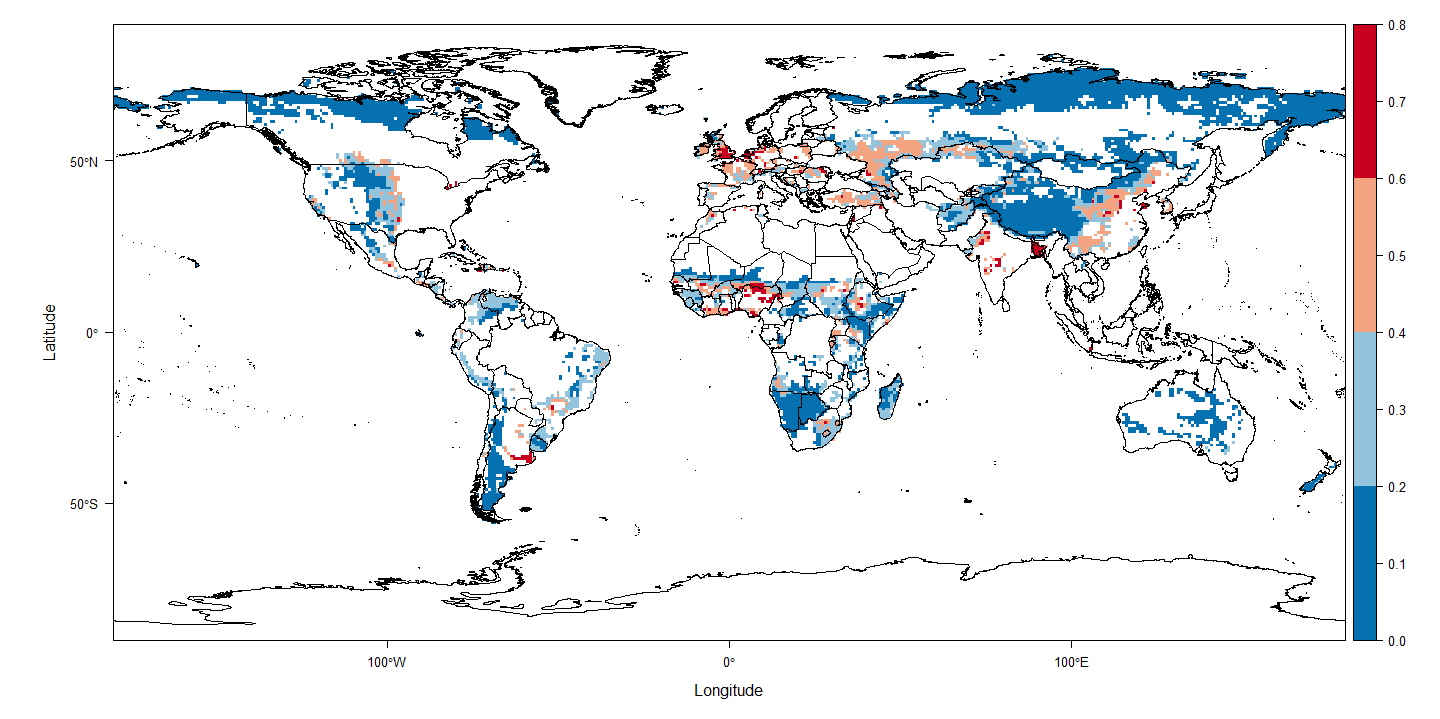

Fig. S1.7. Global human modification of terrestrial systems: grassland. Base map source: ESRI (http:// www. esri. com/ data/ base maps, © Esri, DeLorme Publishing Company).

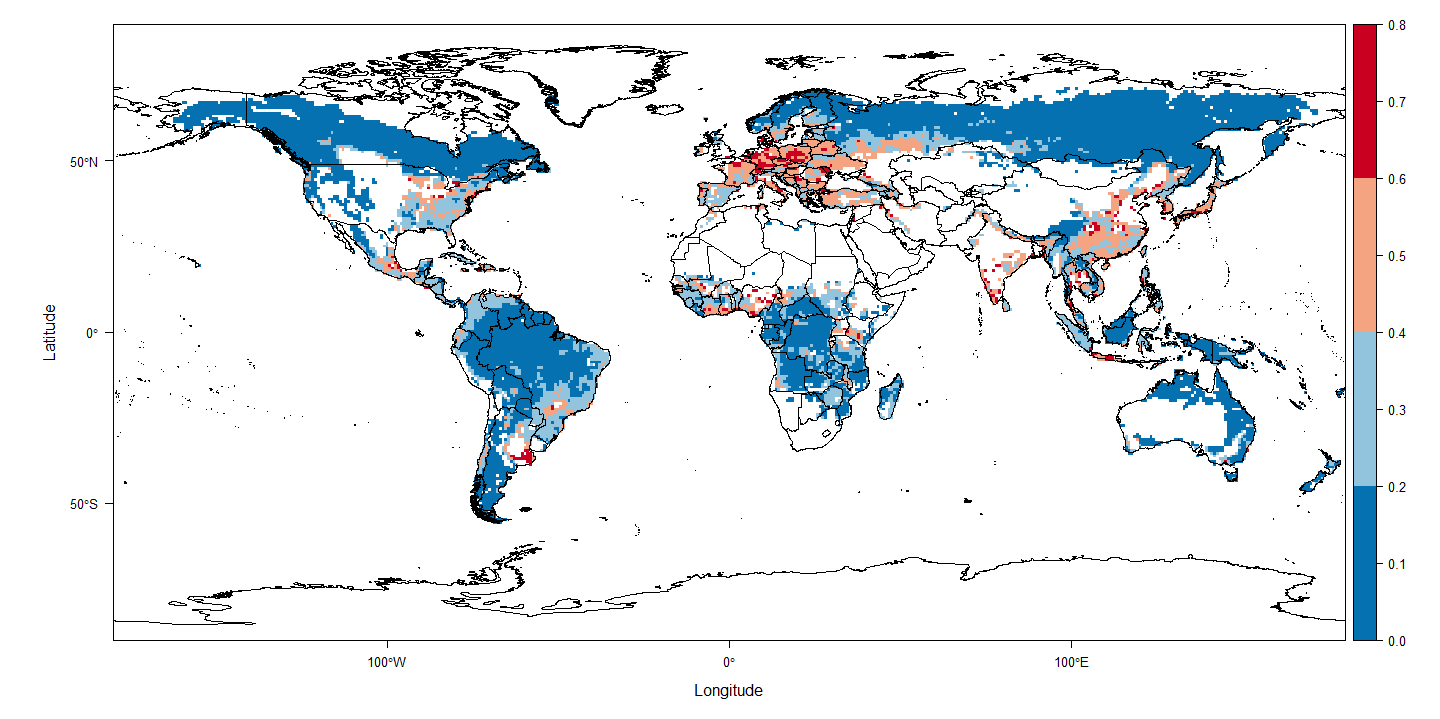

Fig. S1.8. Global human modification of terrestrial systems: forest. Base map source: ESRI (http:// www. esri. com/ data/ base maps, © Esri, DeLorme Publishing Company).

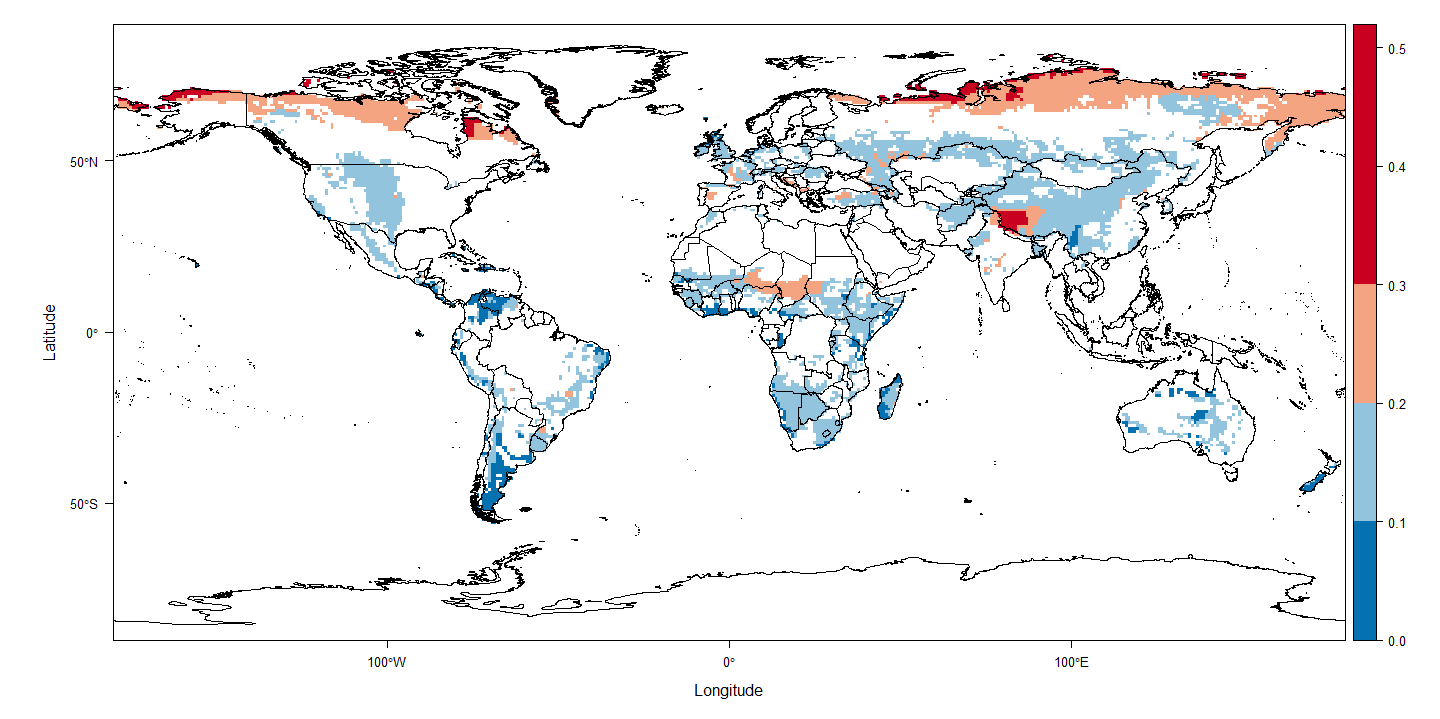

Fig. S1.9. Climate stability from present to 2100: grassland. Base map source: ESRI (http:// www. esri. com/ data/ base maps, © Esri, DeLorme Publishing Company).

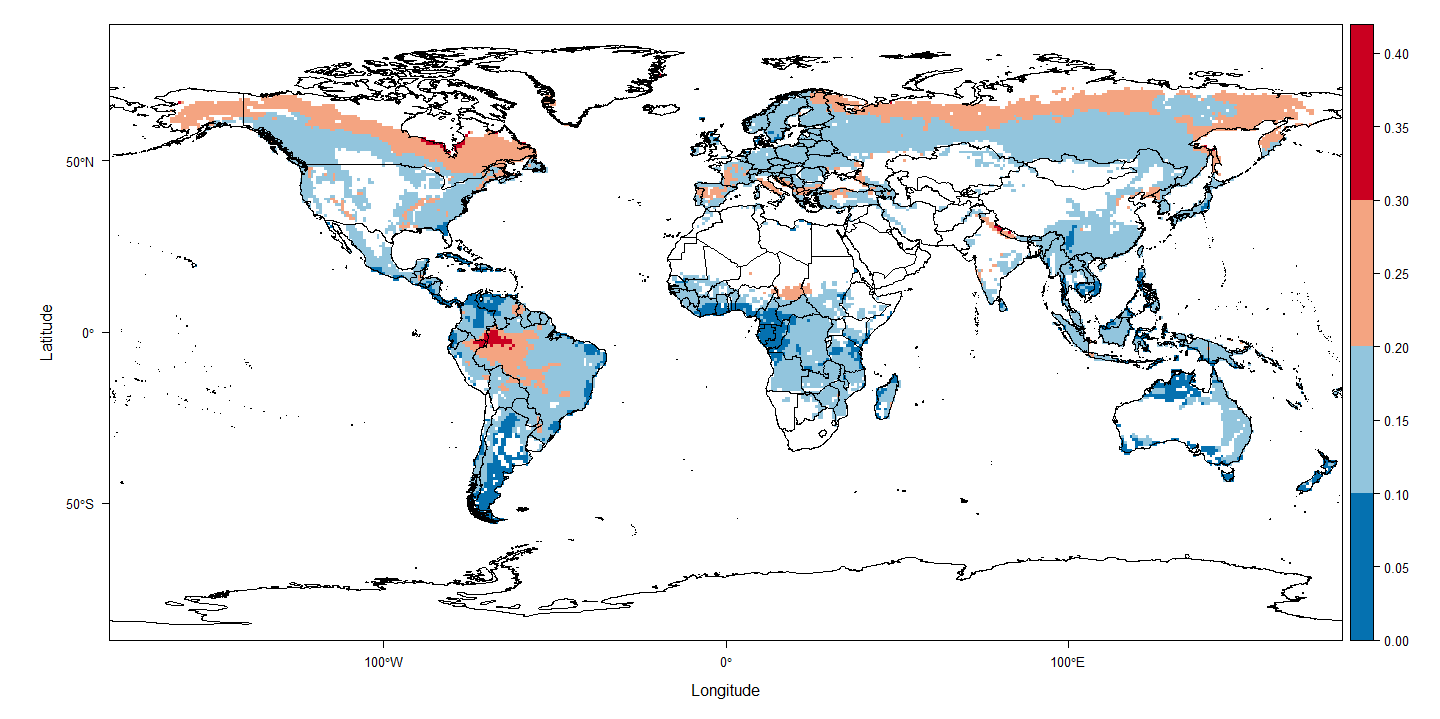

Fig. S1.10. Climate stability from present to 2100: forest. Base map source: ESRI (http:// www. esri. com/ data/ base maps, © Esri, DeLorme Publishing Company).

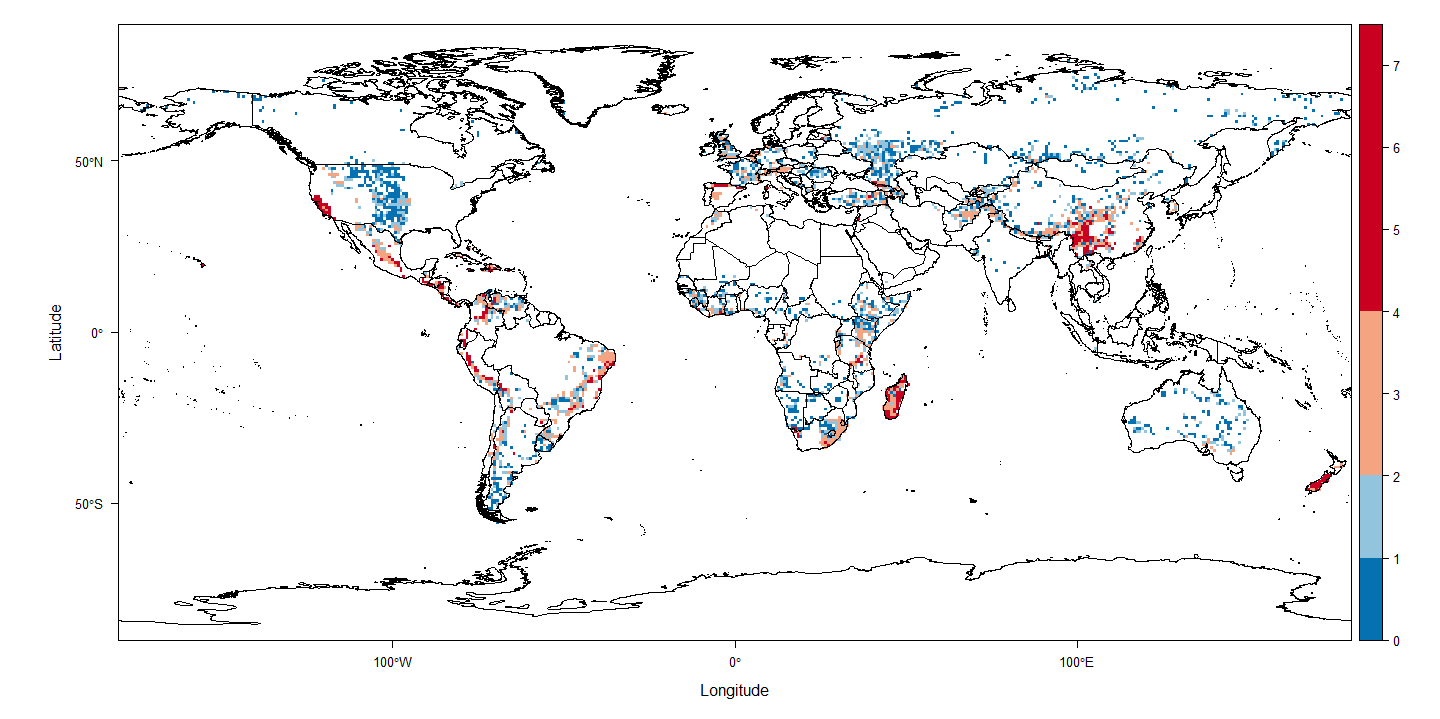

Fig. S1.11. Extinction threat richness (Log10): grassland. Base map source: ESRI (http:// www. esri. com/ data/ base maps, © Esri, DeLorme Publishing Company).

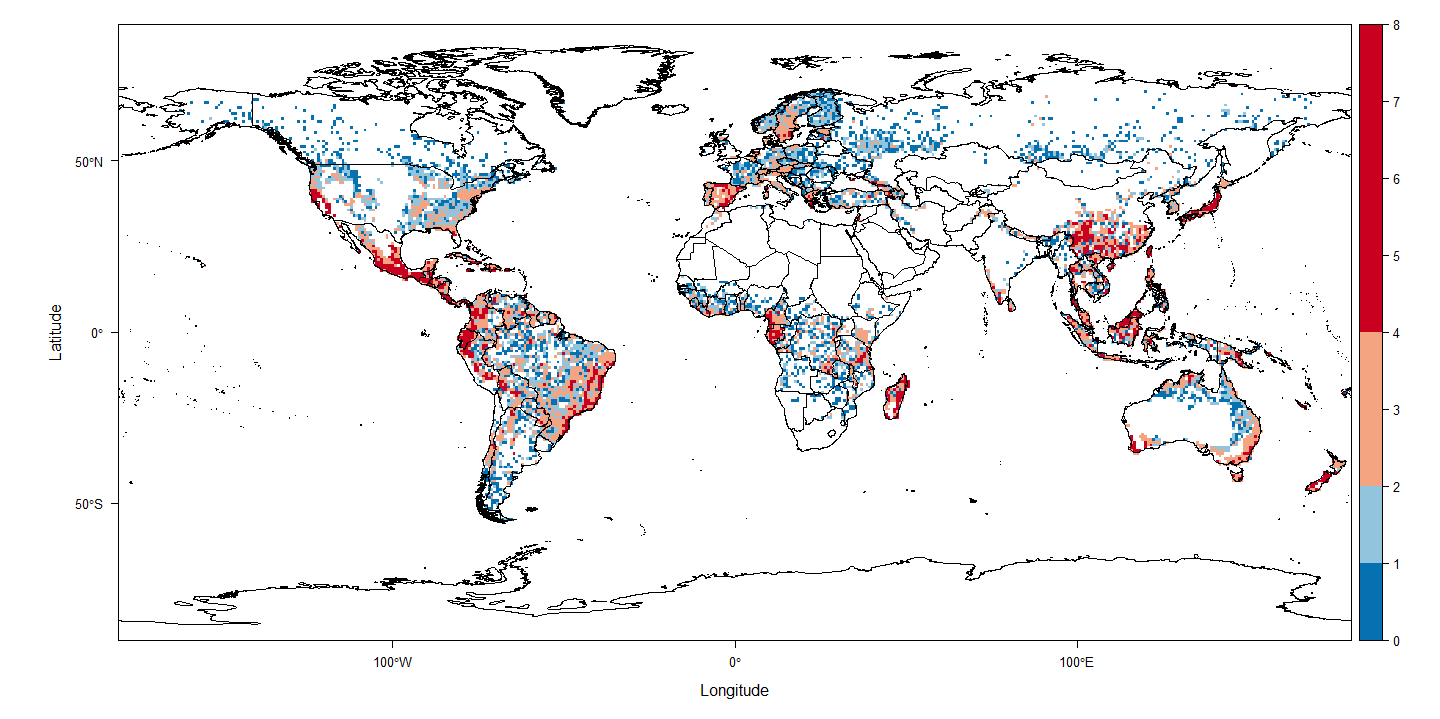

Fig. S1.12. Extinction threat richness (Log10): forest. Base map source: ESRI (http:// www. esri. com/ data/ base maps, © Esri, DeLorme Publishing Company).

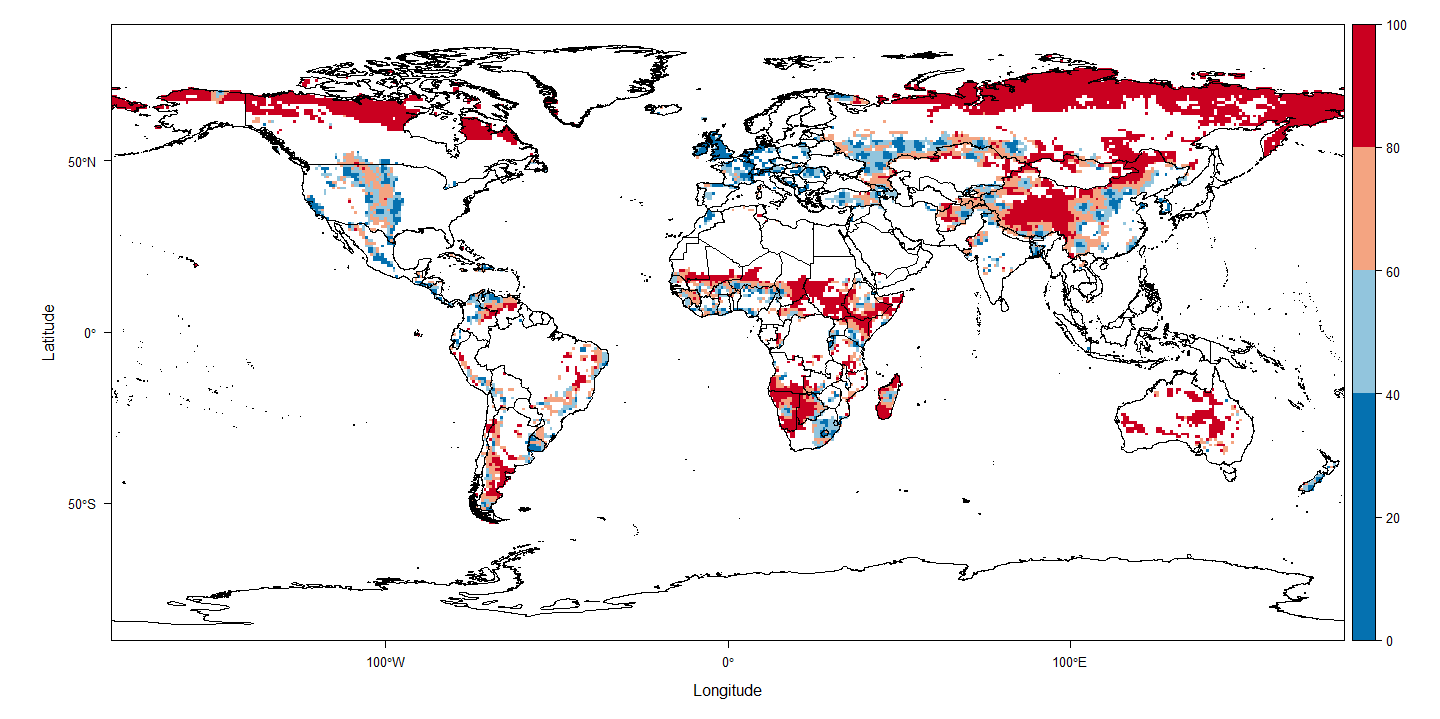

Fig. S1.13. Sampling bias: grassland. Base map source: ESRI (http:// www. esri. com/ data/ base maps, © Esri, DeLorme Publishing Company).

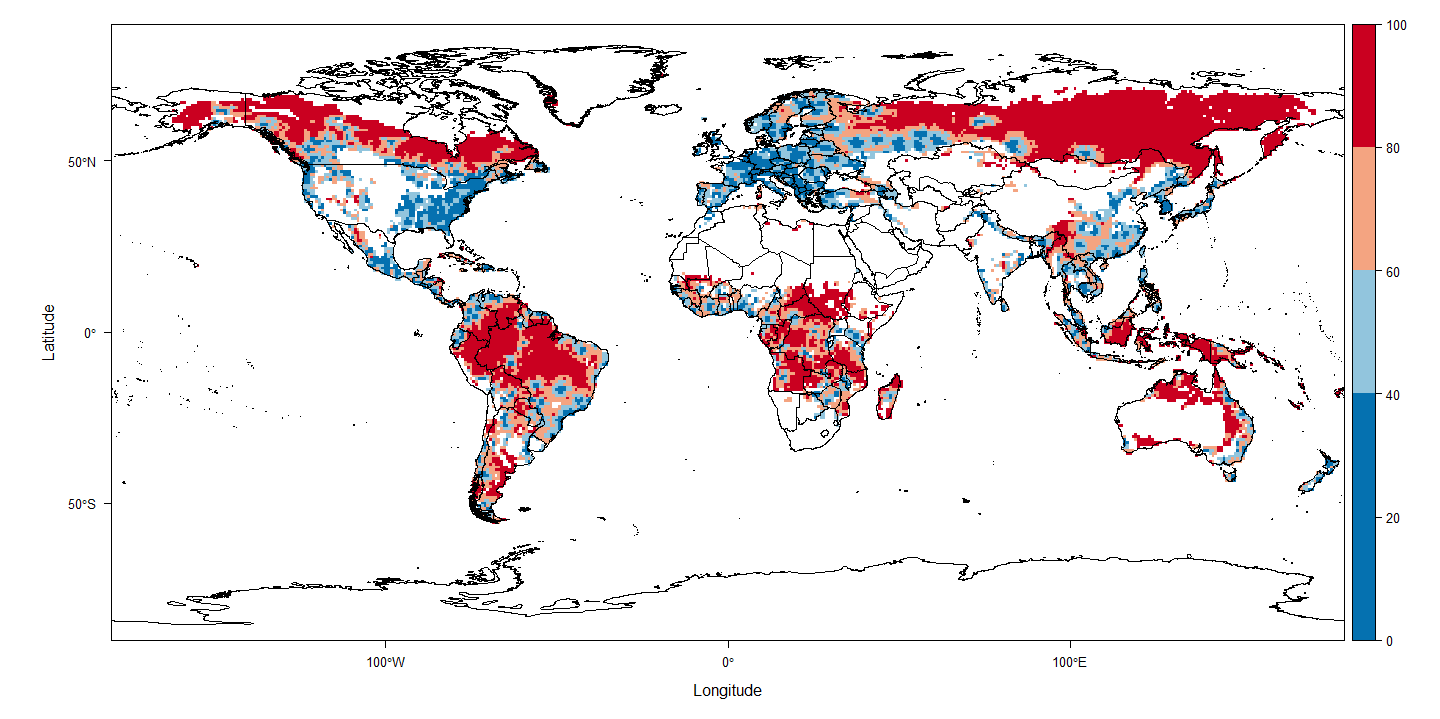

Fig. S1.14. Sampling bias: forest. Base map source: ESRI (http:// www. esri. com/ data/ base maps, © Esri, DeLorme Publishing Company).

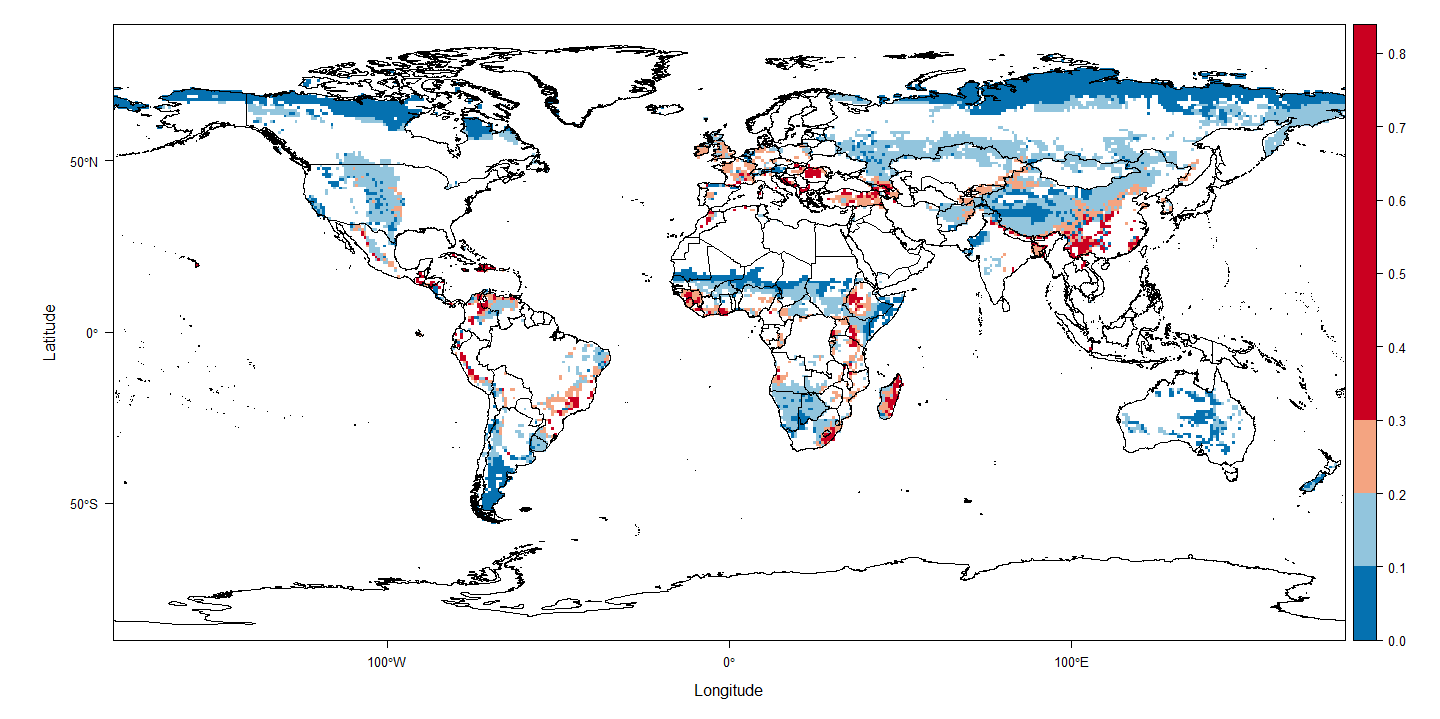

Fig. S1.15. Normalized difference between recorded and expected taxon richness: grassland. Base map source: ESRI (http:// www. esri. com/ data/ base maps, © Esri, DeLorme Publishing Company).

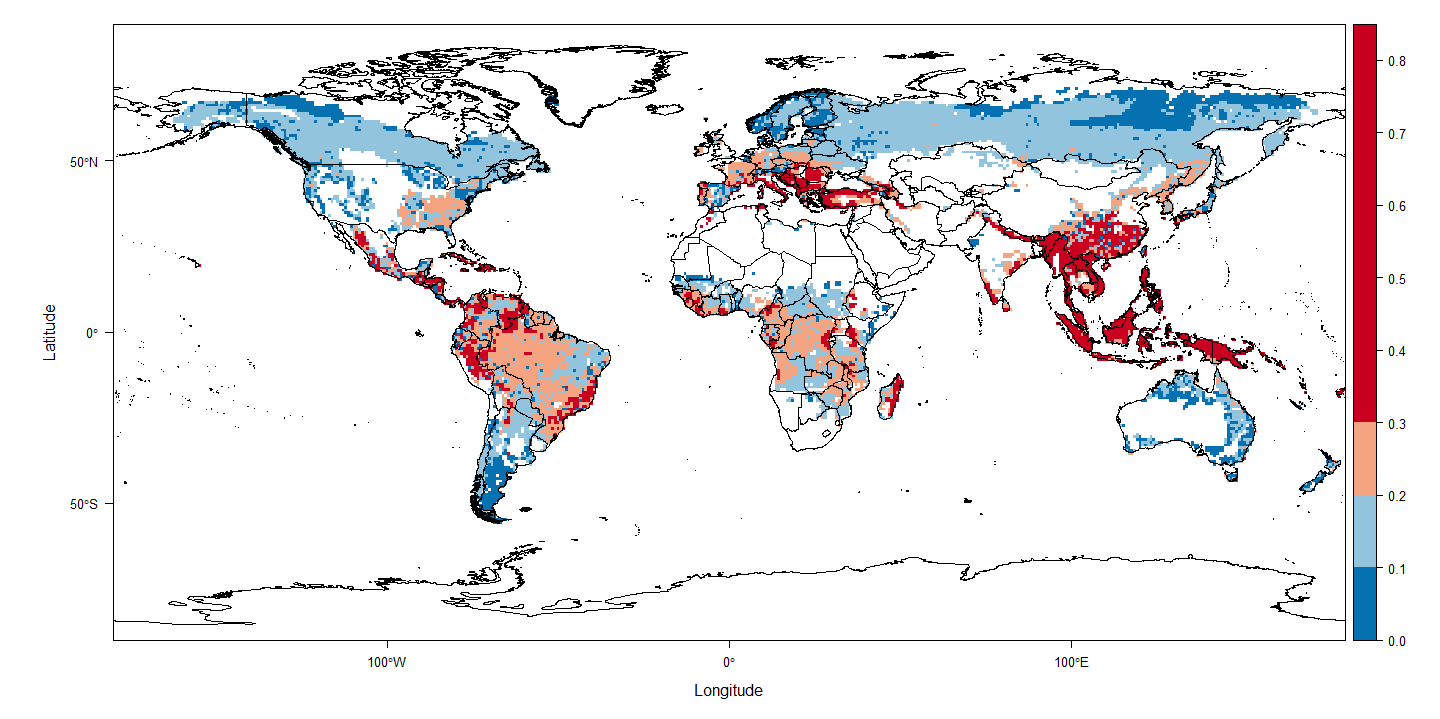

Fig. S1.16. Normalized difference between recorded and expected taxon richness: forest. Base map source: ESRI (http:// www. esri. com/ data/ base maps, © Esri, DeLorme Publishing Company).
