## Supplement 2 for "Setting priorities for the acquisition of primary plant occurrence data"

Supplement 2. TOPSIS model parameters and scores.

Table S2.1. TOPSIS workflow adopted from Roszkowska (2011).

| Step | | Formulae | |
| --- | --- | --- | --- |
| Create an evaluation matrix. A table is created where rows represent alternatives and columns represent criteria. Each cell contains a value representing the performance of an alternative on a specific criterion. | | 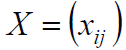 | |
| Normalize the evaluation (decision) matrix to make the criteria comparable, as they might have different scales or units. | | 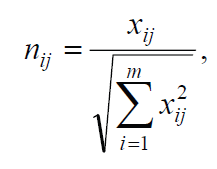 | |
| The normalized values are multiplied by corresponding criteria weights, reflecting the relative importance of each criterion. | | 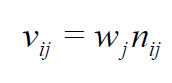 | |
| The best and worst possible values for each criterion are identified. These form the ideal and negative-ideal solutions, respectively. *I* is associated with the criteria having positive impact, and *J*, negative impact. | | | |
| Determine the positive ideal solution. | 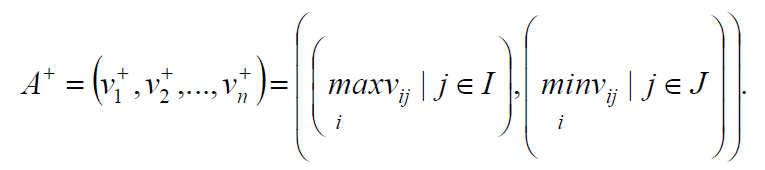 | | |
| Determine the negative ideal solution. | 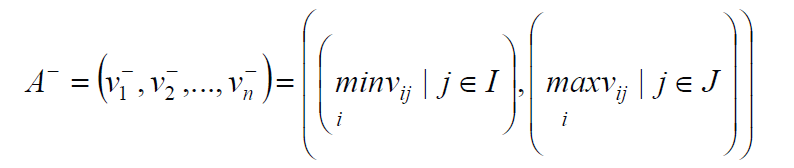 | | |
| The Euclidean distance of each alternative from both the ideal and negative-ideal solutions is calculated. | | | |
| Calculate the distance from the ideal positive solution. | | | 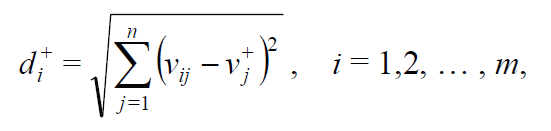 |
| Calculate the distance from the ideal negative solution. | | | 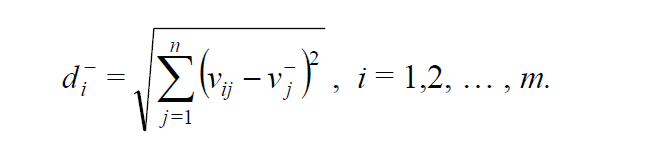 |
| Calculate the relative similarity to the worst solution, and rank the alternatives based on the value of *R.* Alternatives are ranked based on their closeness coefficient, with the highest value indicating the most preferred alternative. | | | 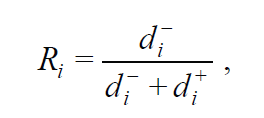 |

Table S2.2. Comparison matrix. For variable names see Table S2.3.

| *bio.carb.water_05* | *csi.past_05* | *rough_05* | *human.modif_05* | *csi.future_05* | *threat.richness_05* | *bias.pc_05* | *gift.sr.interp_05* |
| --- | --- | --- | --- | --- | --- | --- | --- |
| **1.00** | 9.00 | 1.00 | 1.00 | 9.00 | 5.00 | 1.00 | 9.00 |
| 0.11 | **1.00** | 0.11 | 0.11 | 1.00 | 0.20 | 0.11 | 1.00 |
| 1.00 | 9.00 | **1.00** | 1.00 | 9.00 | 5.00 | 1.00 | 9.00 |
| 1.00 | 9.00 | 1.00 | **1.00** | 9.00 | 5.00 | 1.00 | 9.00 |
| 0.11 | 1.00 | 0.11 | 0.11 | **1.00** | 0.20 | 0.11 | 1.00 |
| 0.20 | 5.00 | 0.20 | 0.20 | 5.00 | **1.00** | 0.20 | 5.00 |
| 1.00 | 9.00 | 1.00 | 1.00 | 9.00 | 5.00 | **1.00** | 9.00 |
| 0.11 | 1.00 | 0.11 | 0.11 | 1.00 | 0.20 | 0.11 | **1.00** |

Table S2.3. Weights of selected environmental variables used in TOPSIS models.

| Variable | Variable abbreviated | Benefit | Weight |
| --- | --- | --- | --- |
| Areas of global importance for conserving terrestrial biodiversity, carbon, and water | *bio.carb.water_05* | “ - “ | 0.22 |
| Climate stability from Pliocene (3.3 Ma) to the present | *csi.past_05* | “ - “ | 0.02 |
| Terrain roughness | *rough_05* | “ + “ | 0.22 |
| Global human modification of terrestrial systems | *human.modif_05* | “ - “ | 0.22 |
| Climate stability from present to 2100 | *csi.future_05* | “ + “ | 0.02 |
| Extinction threat | *threat.richness_05* | “ + “ | 0.07 |
| Sampling bias | *bias.pc_05* | “ + “ | 0.22 |
| Normalized difference between recorded and expected taxon richness | *gift.sr.interp_05* | “ + “ | 0.02 |

Table S2.4. TOPSIS model extents and projections.

| Region | Extent (min LON, max LON, min LAT, max LAT) | Biome | Projection |
| --- | --- | --- | --- |
| Global | (-180, 180, -90, 90) | Grassland | Eckert IV |
| Global | (-180, 180, -90, 90) | Forest | Eckert IV |
| South America | (-81, -35, -55, 12) | Grassland | Transverse cylindrical equal area |
| South America | (-81, -35, -55, 12) | Forest | Transverse cylindrical equal area |
| Africa | (-26, 60, -42, 37) | Grassland | Equatorial Lambert azimuthal equal area |
| Africa | (-26, 60, -42, 37) | Forest | Equatorial Lambert azimuthal equal area |
| Southeast Asia | (65, 160, -10, 50) | Grassland | Albers equal-area conic |
| Southeast Asia | (65, 160, -10, 50) | Forest | Albers equal-area conic |
| Siberia and Russian Far East | (50, 180,40, 76) | Grassland | Albers equal-area conic |
| Siberia and Russian Far East | (50, 180,40, 76) | Forest | Albers equal-area conic |
| Central and North America | (-165, -50,5, 70) | Grassland | Eckert IV |
| Central and North America | (-165, -50,5, 70) | Forest | Eckert IV |

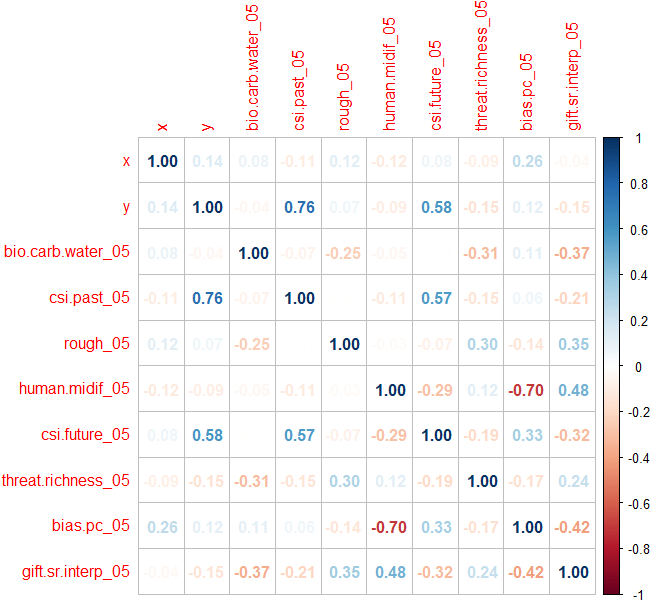

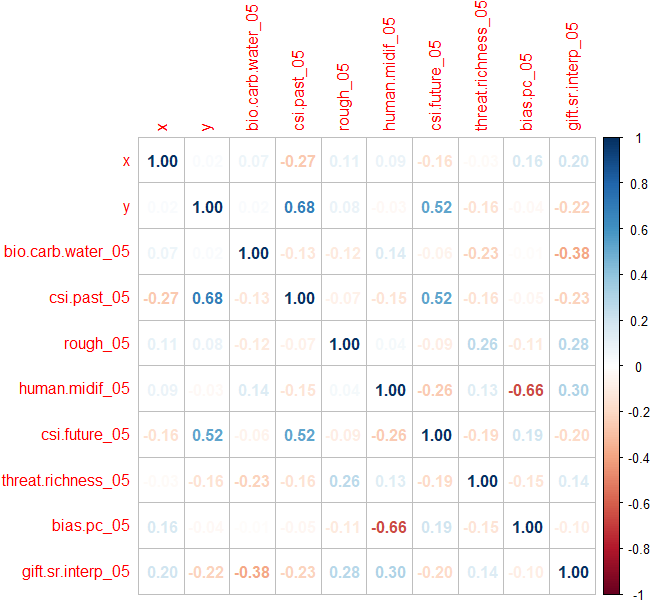

Fig. S2.1. Variable correlations within the global extent limited by grassland (top) and forest (bottom) cover.

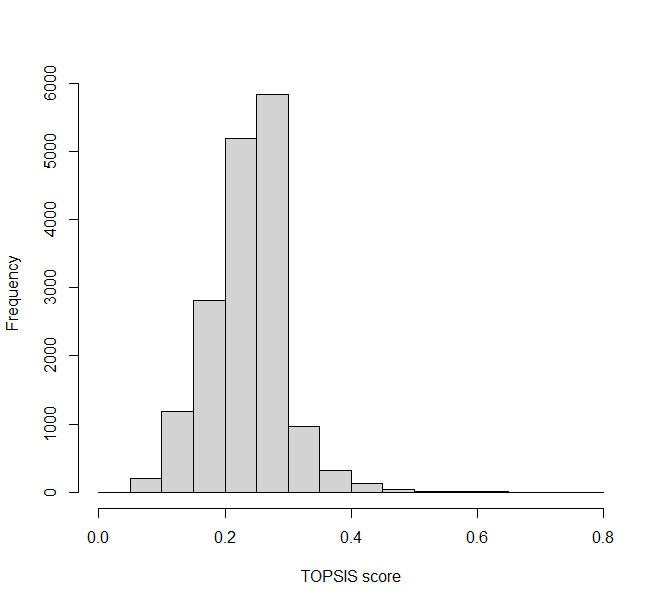

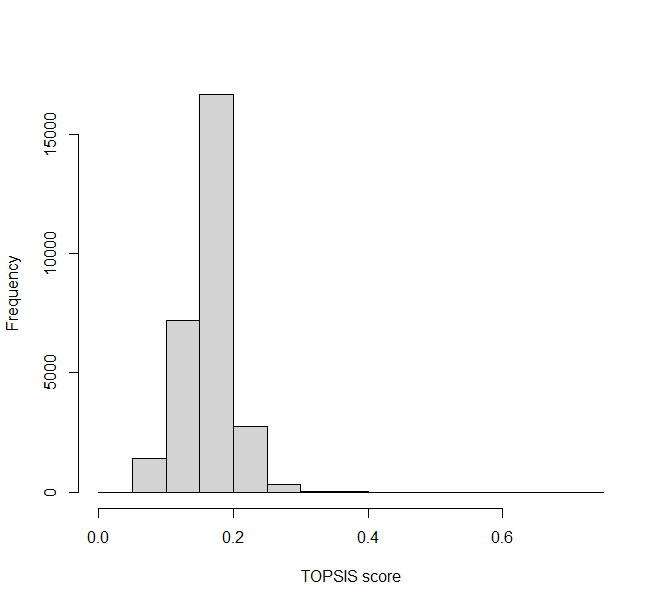

Fig. S2.2. Histograms of TOPSIS scores for the global extent: grassland (top) and forest (bottom).

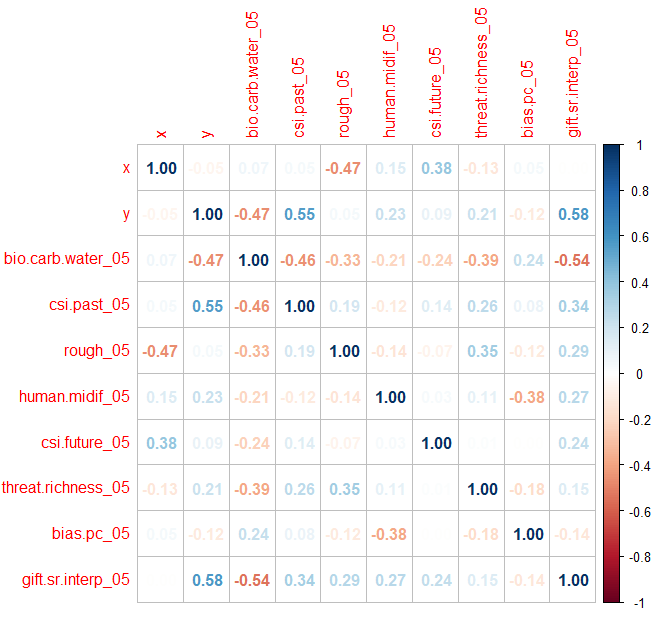

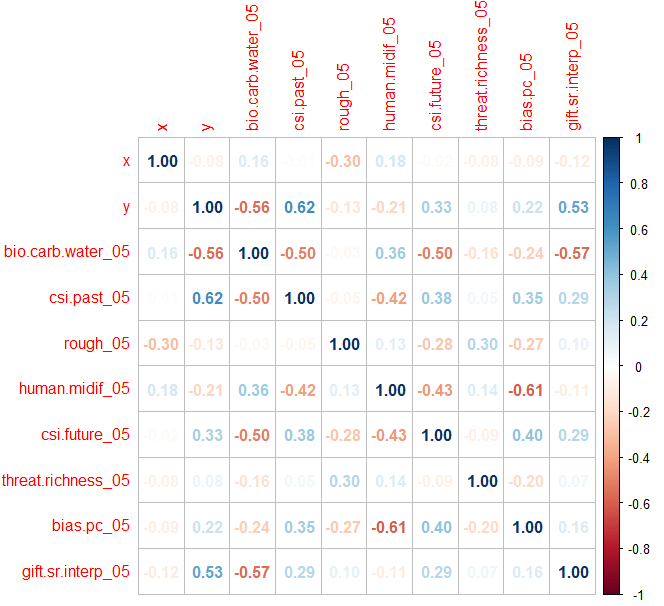

Fig. S2.3. Variable correlations within the South America extent limited by grassland (top) and forest (bottom) cover.

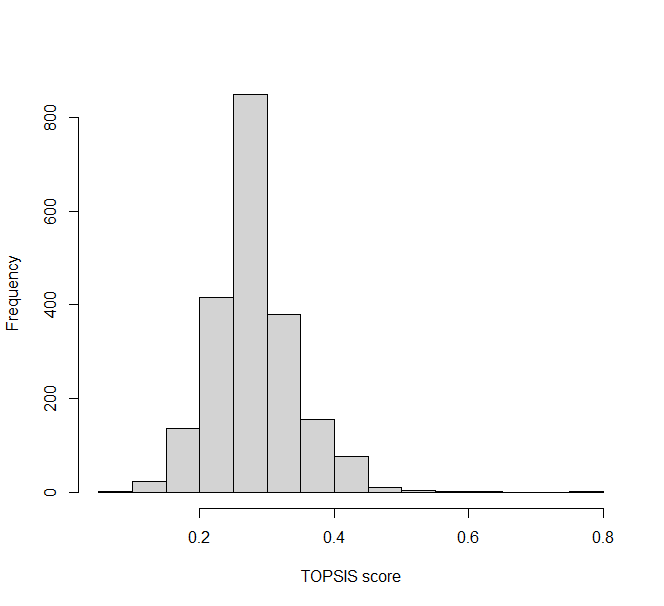

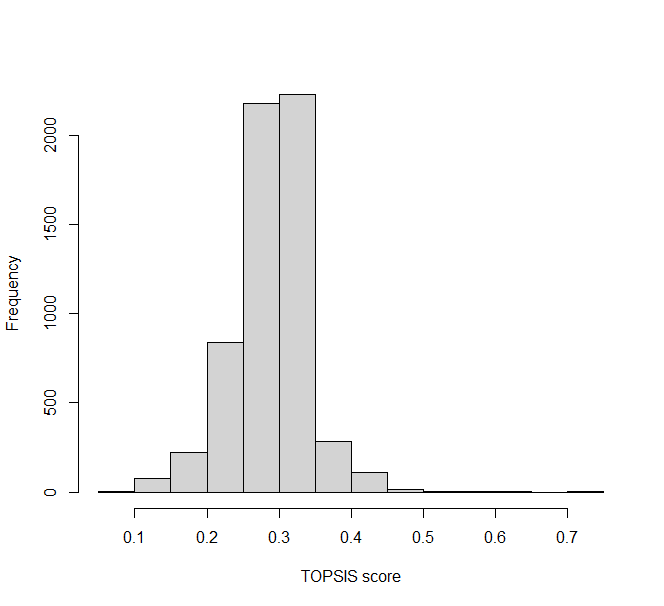

Fig. S2.2. Histograms of TOPSIS scores for South America extent: grassland (top) and forest (bottom).

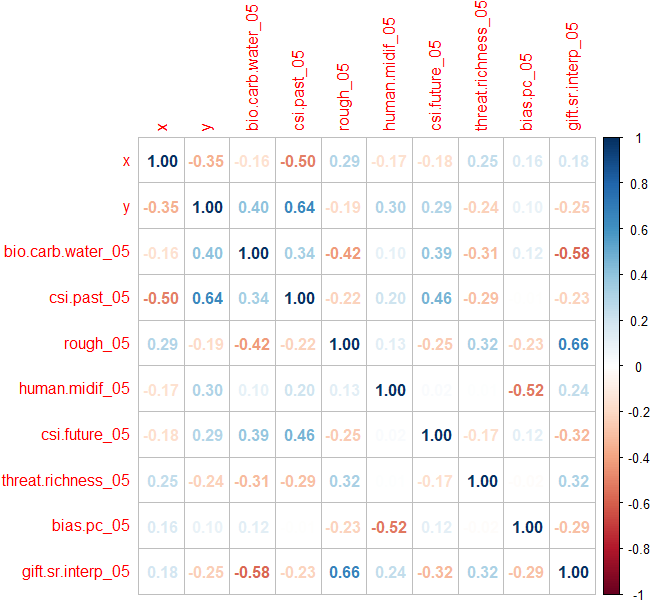

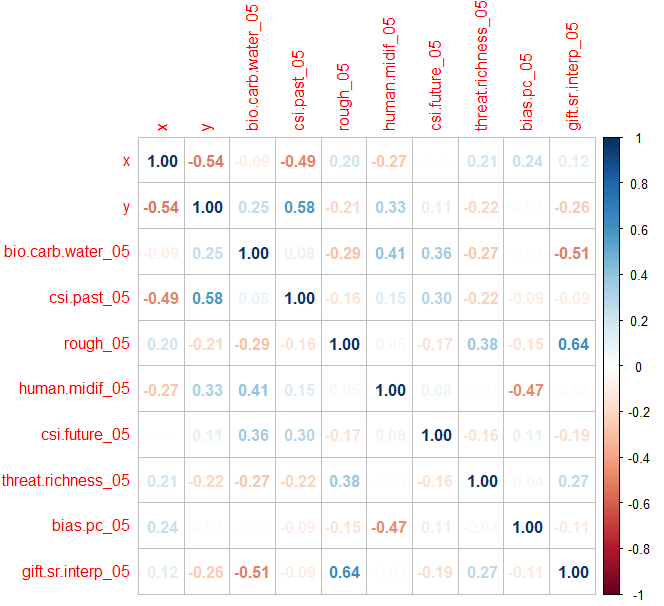

Fig. S2.3. Variable correlations within the African and Madagascan extent limited by grassland (top) and forest (bottom) cover.

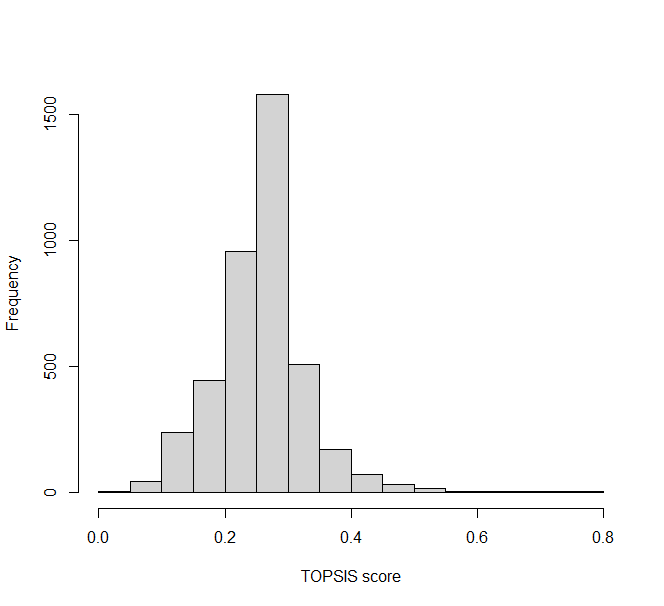

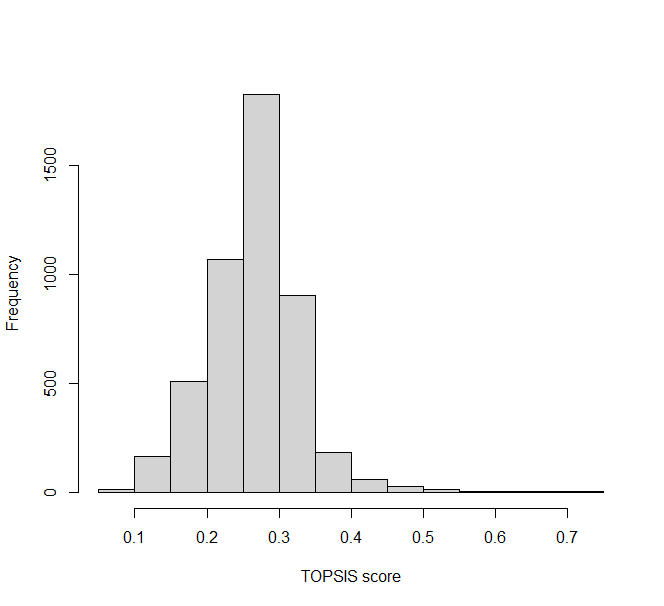

Fig. S2.4. Histograms of TOPSIS scores for the African and Madagascan extent limited by grassland (top) and forest (bottom) cover.

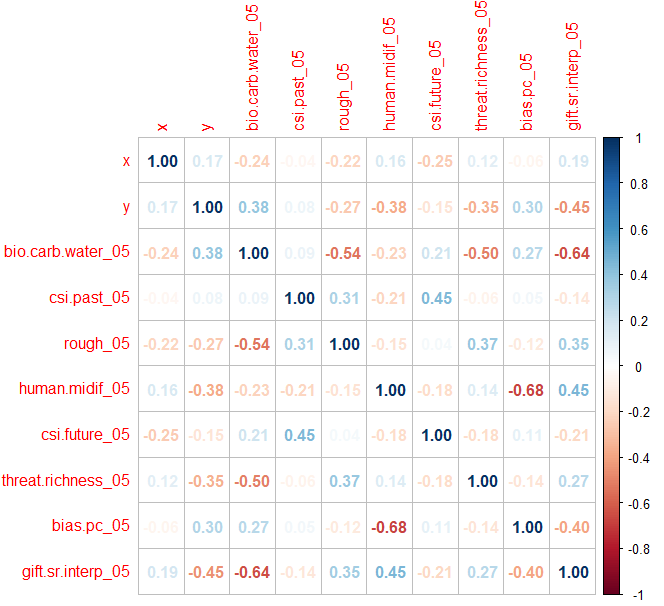

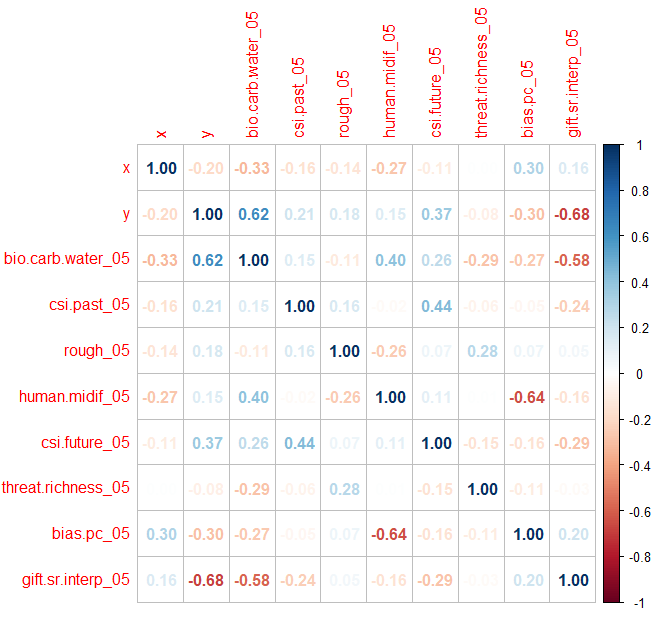

Fig. S2.5. Variable correlations within the Southeast Asia extent limited by grassland (top) and forest (bottom) cover.

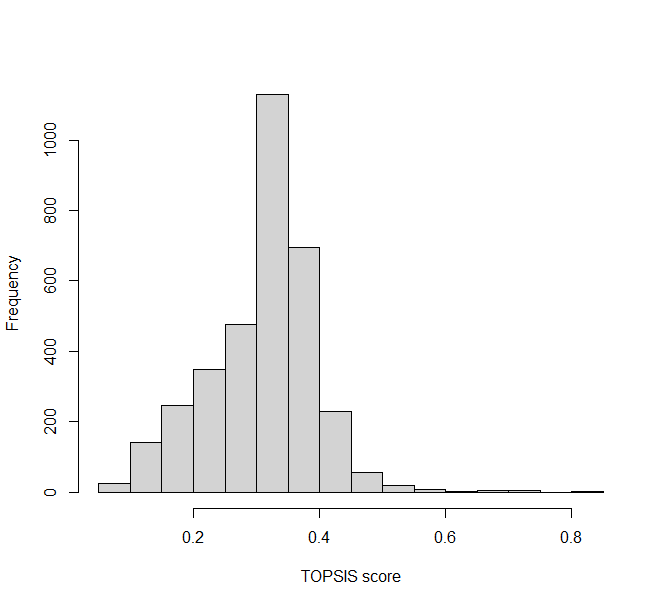

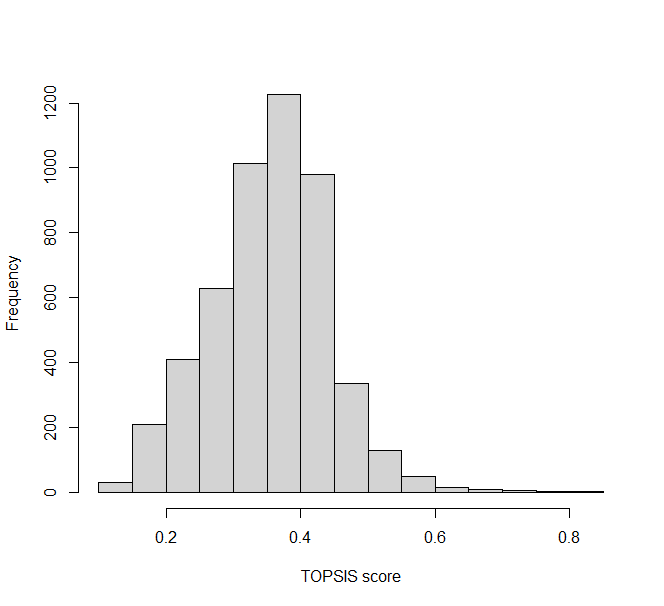

Fig. S2.6. Histograms of TOPSIS scores for the Southeast Asia extent limited by grassland (top) and forest (bottom) cover.

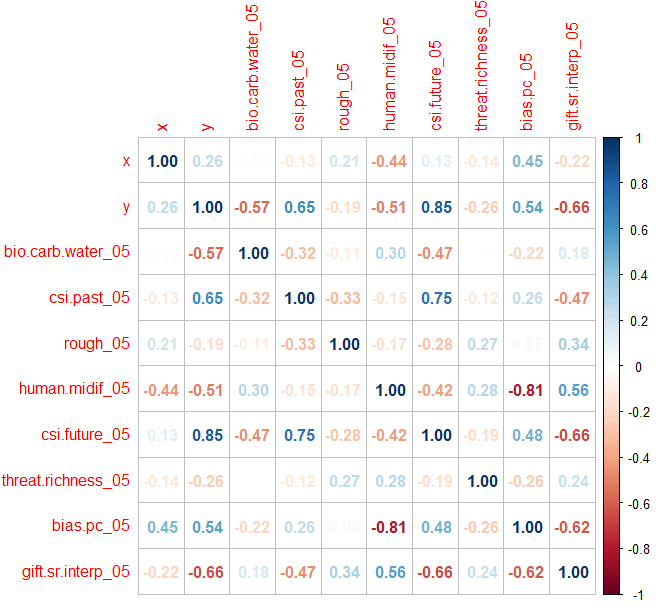

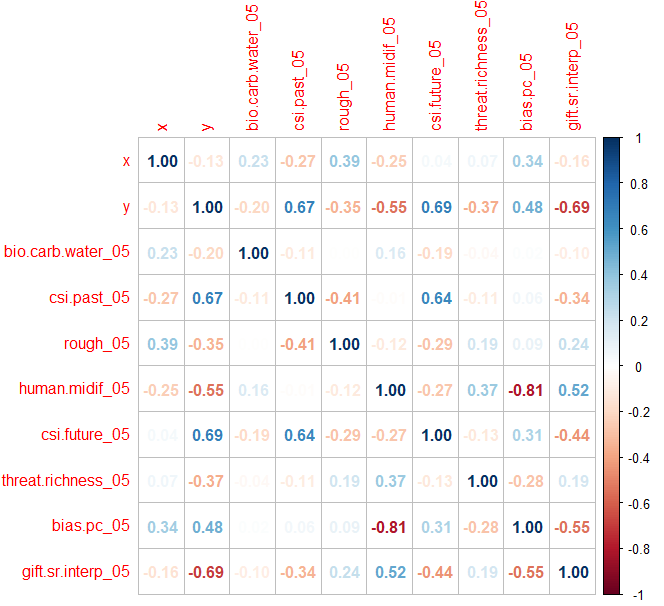

Fig. S2.7. Variable correlations within the Siberia and Russian Far East extent limited by grassland (top) and forest (bottom) cover.

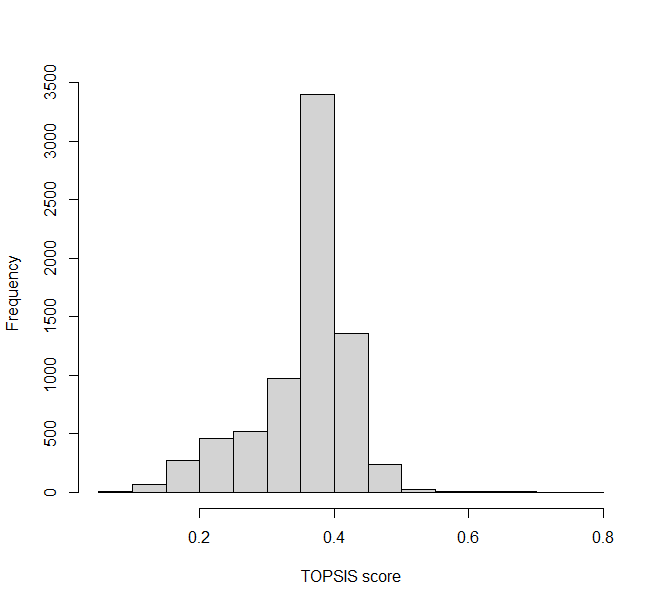

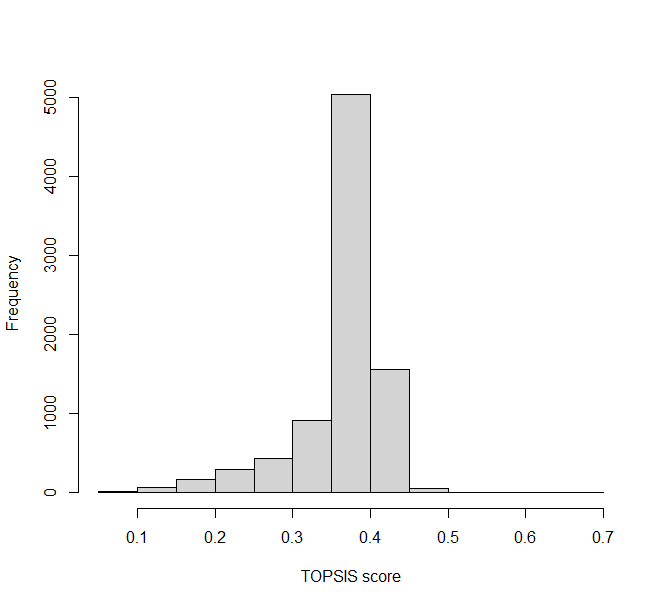

Fig. S2.8. Histograms of TOPSIS scores for the Sibiria and Russian Far East extent limited by grassland (top) and forest (bottom) cover.

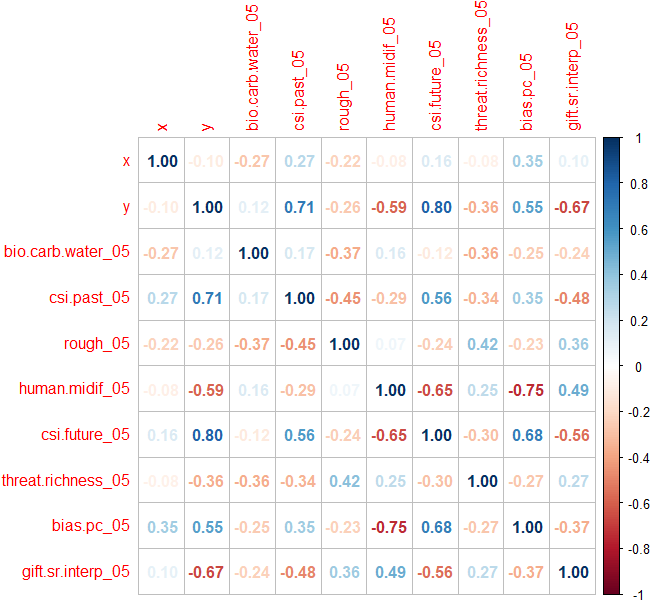

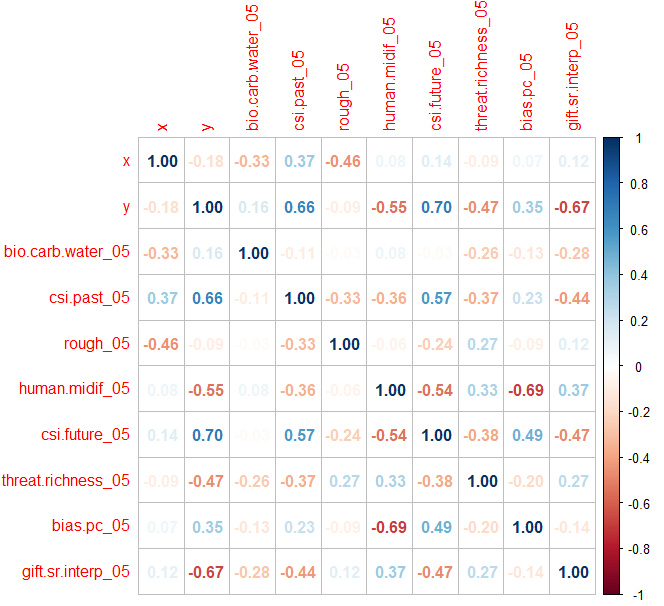

Fig. S2.9. Variable correlations within the Central and North America extent limited by grassland (top) and forest (bottom) cover.

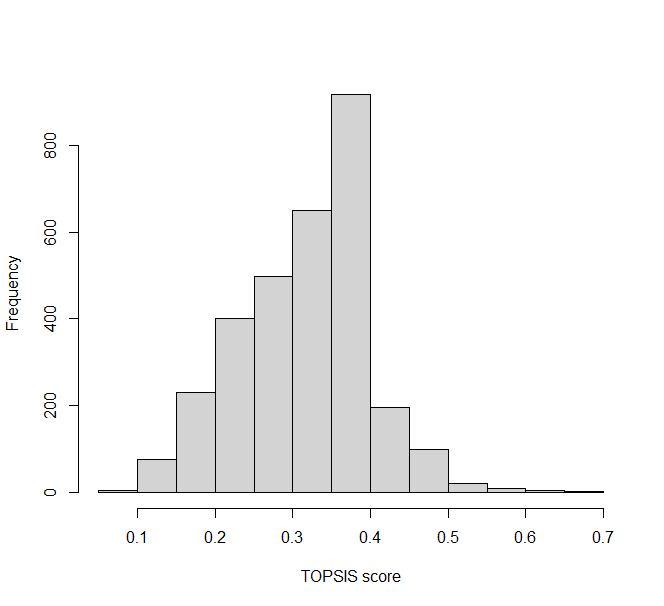

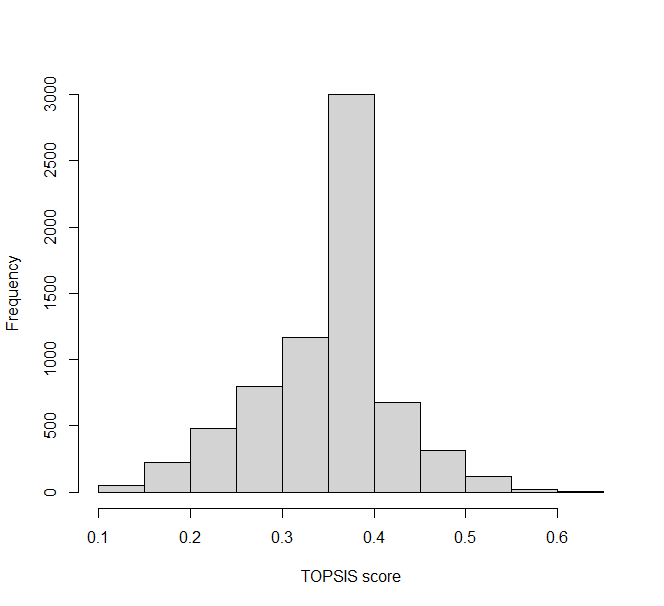

Fig. S2.10. Histograms of TOPSIS scores for the Central and North America extent limited by grassland (top) and forest (bottom) cover.
