## Supplement 4 for "Setting priorities for the acquisition of primary plant occurrence data"

Table 4.1. Summary of proportion of consensus.

| Region | Supported | Not Supported | Consensus Proportion |
| --- | --- | --- | --- |
| North & Meso- America | 11 | 0 | 100.00% |
| Africa & Madagascar | 53 | 3 | 95.00% |
| Siberia & Russian Far East | 4 | 0 | 100.00% |
| South America | 24 | 2 | 92.00% |
| South Asia, Southeast Asia, East Asia | 23 | 4 | 85.00% |

Table 4.2. Field expert assessment.

| Analysis | Region | Fig. no. | Areas of highest Priority areas (top 25%) confirmed | Highest priority vegetation class | Field expert reviewers | Areas not supported by reviewers |
| --- | --- | --- | --- | --- | --- | --- |
| Global, regional | Africa & Madagascar | 2,3 | Congolian | forest, non-forest dominated | Martin Cheek (MC), Carol Jongkind (CJ) | SW Ghana, Ivory Coast, Liberia: white sand coastal thickets should be high priority (**MC, CJ**). Liberia: naturally open rocky areas (1% or 2% of the total surface) should be high priority as very poorly sampled (**CJ**). High priority areas in African (Guineo-Congolian Africa) are better mapped, more of Liberia should be red and not pink or pale blue. Sahel, southern Chad, Mali, much of Senegal: It appears that open woodland has been misclassified as forest. A large block of this area mapped as low priority forest comprises secondary vegetation and not forest (0.5 to 20%), confusingly this intergrades with non-forest dominated land to the north (which has never been forested) (**MC**). |
| Global, regional | Africa & Madagascar | 2,3, 4 | Zambeziana | forest, non-forest dominated | Iain Darbyshire (ID), MC, CJ |  |
| Regional | Africa & Madagascar | 2,3, 4 | Ethiopian | forest, non-forest dominated | Sebsebe Demissew (SD), Ib Friis (IB) | Forested: this does not agree with the observed reality in Ethiopia. SE Ethiopia which is shown in white and blue is given a low priority for further data gathering but should be high priority (**SD**). Non-forest dominated *vs* forest dominated distinction is not really appropriate to use to the east of the Ethiopian highlands because almost all of that area is covered with deciduous bushland. Areas in the south-western part of Ethiopia indicated as low priority in Fig. 4 should be given a much higher priority as they have the highest number of endemics. I think the Ethiopian highlands are ranked as too high apriority, most of the highlands used to be dry forest but are now farmland and secondary grassland, conversely the areas of the eastern lowlands and Somalia ranked as too low a priority (**IB**). |
| Global, regional | Africa & Madagascar | 2,3, 4 | Southern Africa | non-forest dominated | MC, CJ |  |
| Global, regional | Africa & Madagascar | 2,3, 5 | Madagascar | forest, non-forest dominated | MC, CJ, Justin Moat (JM), Bat Maria Vorontsova (BV) | East coast non-forest dominated are probably very secondary and just removed forest (**JM**). There are four locations on the east coast where non-forest dominated areas is marked red. This is secondary forest and not a high priority for data acquisition (**MC**). |
| Global |  | 2, 3 | Dolomites | forest, non-forest |  |  |
| Global |  | 2, 3 | Caucasus | forest, non-forest |  |  |
| Global, regional | Siberia and Russian Far East | 2,3, 5 | Siberia | forest, non-forest | Nadia Bystriakova (NB), Andrey Erst (AE) |  |
| Global, regional | Siberia and Russian Far East | 2,3, 5 | Altai | forest, non-forest | Nadia Bystriakova (NB), Andrey Erst (AE) |  |
| Global, regional | Siberia and Russian Far East | 2,3, 5 | Kamchatka | forest, non-forest | Nadia Bystriakova (NB), Andrey Erst (AE) |  |
| Global, regional | Siberia and Russian Far East | 2,3, 5 | Ural Mountains | forest, non-forest | Nadia Bystriakova (NB), Andrey Erst (AE) |  |
| Global, regional | E Asia | 2, 3, 4 | Himalaya Mountains | forest, non-forest dominated | Rajib Gogoi (RG) |  |
| Global, regional | E Asia | 2, 3, 4 | Tibetan / Xizang Plateau | forest |  |  |
| Global, regional | E Asia | 2, 3, 4 | Guizhou – Yunnan plateau | forest | Long-Fei Fu (LF), Yi-Gang Wei (YG) |  |
| Global, regional | E Asia | 2, 3, 4 | Sunda shelf | forest | Gemma Bramley (GB), Laura Jennings (LJ), Himmah Rustiami (HR), Liam Trethowan (LT),Tim Utteridge (TU) | Indonesia, Borneo: Timor is the most diverse of all the islands and I would not expect it to be blue. Colours on Borneo should be flipped north to south: Sabah, Sarawak and Brunei are much better surveyed than Kalimantan. (**LJ)** |
| Global, regional | E Asia | 2, 3, 4 | Wallacea | forest | GB, LT, TU |  |
| Global, regional | E Asia | 2, 3, 4 | Sahul shelf | forest | GB, LT, TU | New Guinea: Southern savannas are less diverse though very under sampled, so would expect medium priority here (**LJ**). The blue pixel in north PNG are odd: I would expect any low priority areas to be close to Lae or Port Moresby. I can't think of any mass sampling efforts in the Sepik: maybe a centroid problem? Mount Jaya appears over-priroitised (**LT**). |
| Global, regional | E Asia | 2, 3, 4 | Tian Shan | non-forest dominated |  |  |
| Global, regional | E Asia | 2, 3, 4 | Hindu Kush | non-forest dominated |  |  |
| Global, regional | E Asia | 2, 3, 4 | Karakorum | non-forest dominated |  |  |
| Global, regional | South America | 2, 3, 6 | Andes | forest, non-forest dominated land | JM, Carlos Reynel (CR) | Peru: odd patch in dry lands, South Lima to Paracus. I wouldn't classify as forest JM |
| Global, regional | South America | 2, 3, 6 | Amazonia | forest |  |  |
| Global, regional | South America | 2, 3, 6 | Atlantic Forest | forest |  |  |
| Global, regional | South America | 2, 3, 6 | Caatinga | forest | Eimear Nic Lughadha (ENL) | Brazil: Analysis might have (slightly) over prioritised the cerrado (also poorly sampled) in the SW of Piaui rather than the Caatinga. Cerrado species tend to be more widespread than Caatinga species (**ENL**). |
| Global, regional | South America | 2, 3, 6 | Cerrado | forest, non-forest dominated land | Gloria Céspedes (GC), Juana de Egea (JdE), ENL | Paraguay: area marked as non-forest dominated corresponds mainly to deforested areas. E.g. Eastern Paraguay, Cordillera de los Altos and the forests of Caaguazú. Western region of the country, the non-forest dominated land areas correspond to pastures implanted by the Mennonites; this area is completely degraded due to the high agricultural productivity of the area (**GC, JdE**). |
| Global, regional | South America | 2, 3, 6 | Chiqitania | forest | Bente Klitgaard (BK), Carla Malodonado (CM), Alfredo Fuentes (AF) |  |
| Global, regional | South America | 2, 3, 6 | Pampas del Beni | non-forest dominated land | Bente Klitgaard (BK), Carla Malodonado (CM), Maira Martínez (MM), Alfredo Fuentes (AF) |  |
| Global, regional | South America | 2, 3, 6 | Choco | forest-dominated | Sebastian Tello (ST) | Parts of the Ecuadorin Chocó are identified as not high priority should be high priority (**ST**) |
| Global, regional | South America | 2, 3, 6 | Pampas | non-forest dominated land | ENL | Brazil: southern non-forest dominated lands (part of the Pampa biome), the area furthest to the south, bordering Uruguay, is absent from the map but should be recognised as high priority, including highly range-restricted species also found just over the border in north (ENL). |
| Global, regional | North & Meso- America | 2, 3, 7 | Mesoamerica | forest | Alexandre Monro (AM), Daniel Santamaría Aguilar (DSA) | Guatemala, El Salvador: coastal forest should be high priority due to combination of threat and low sample effort. |
| Global, regional | North & Meso- America | 2, 3, 7 | Sierra Madre | non-forest dominatedland |  |  |
| Global, regional | North & Meso- America | 2, 3, 7 | Taiga | forest |  |  |
| Global, regional | North & Meso- America | 2, 3, 7 | Hudson Plain | forest |  |  |
