## Supplementary figures and images for "Setting priorities for the acquisition of primary plant occurrence data"

### Supplement 5

**Supplement 5.** Technique for Order of Preference by Similarity to Ideal Solution (TOPSIS)

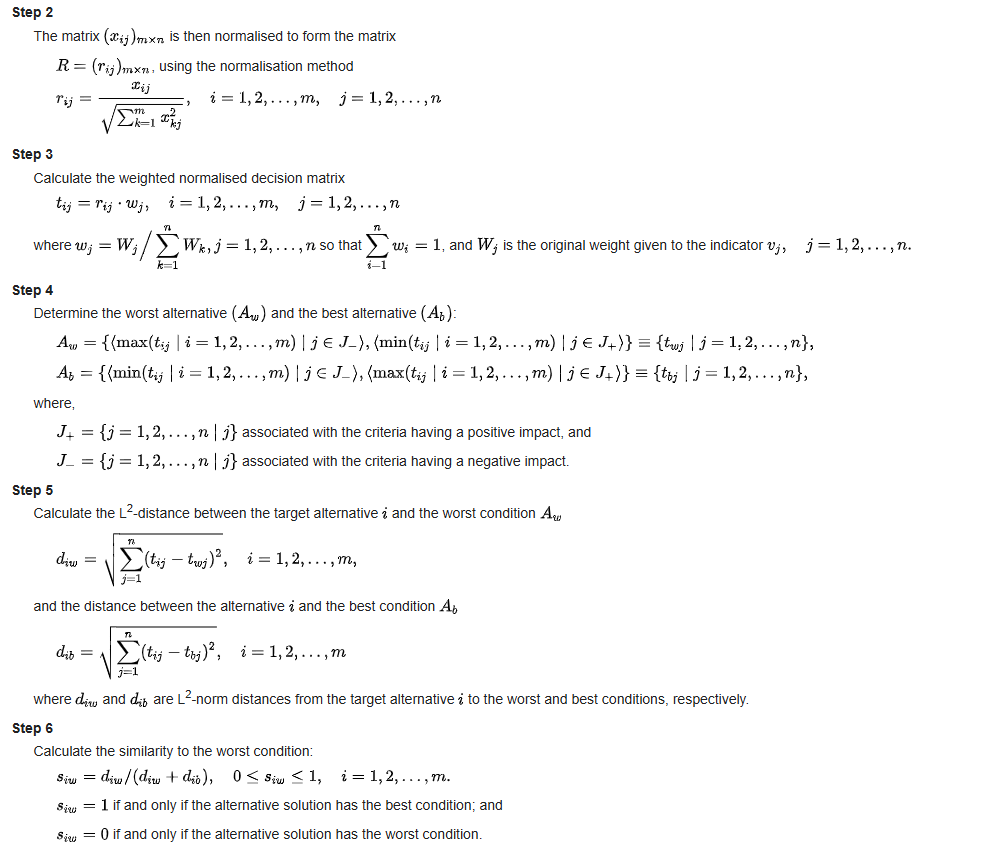

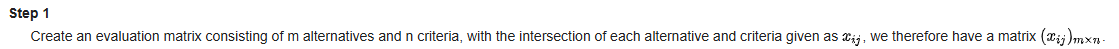

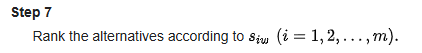
